# Efficient multiplex non-viral engineering and expansion of polyclonal γδ CAR-T cells for immunotherapy

**DOI:** 10.1101/2024.09.03.611042

**Authors:** Jacob Bridge, Matthew J. Johnson, Jihyun Kim, Sophia Wenthe, Joshua Krueger, Bryce Wick, Mitchell Kluesner, Andrew T. Crane, Jason Bell, Joseph G. Skeate, Branden S. Moriarity, Beau R. Webber

**Affiliations:** Department of Pediatrics, University of Minnesota, Minneapolis, MN, USA; Masonic Cancer Center, University of Minnesota, Minneapolis, MN, USA; Center for Genome Engineering, University of Minnesota, Minneapolis, MN, USA; Department of Genetics, Cell Biology and Development, University of Minnesota, Minneapolis, MN, USA; Molecular and Cellular Biology Graduate Program, University of Washington, Seattle, WA, USA; Human Biology Division, Fred Hutchinson Cancer Research Center, Seattle, WA, USA; Stem Cell Institute, University of Minnesota, Minneapolis, MN, USA

**Author notes:** These authors contributed equally to this work (Jacob Bridge, Matthew J. Johnson). These authors co-lead this work (Branden S. Moriarity, Beau R. Webber).

## Abstract

Gamma delta (γδ) T cells are defined by their unique ability to recognize a limited repertoire of non-peptide, non-MHC-associated antigens on transformed and pathogen-infected cells. In addition to their lack of alloreactivity, γδ T cells exhibit properties distinct from other lymphocyte subsets, prompting significant interest in their development as an off-the-shelf cellular immunotherapeutic. However, their low abundance in circulation, heterogeneity, limited methods for *ex vivo* expansion, and under-developed methodologies for genetic modification have hindered basic study and clinical application of γδ T cells. Here, we implement a feeder-free, scalable approach for *ex vivo* manufacture of polyclonal, non-virally modified, gene edited chimeric antigen receptor (CAR)-γδ T cells in support of therapeutic application. Engineered CAR-γδ T cells demonstrate high function *in vitro* and and *in vivo.* Longitudinal *in vivo* pharmacokinetic profiling of adoptively transferred polyclonal CAR-γδ T cells uncover subset-specific responses to IL-15 cytokine armoring and multiplex base editing. Our results present a robust platform for genetic modification of polyclonal CAR-γδ T cells and present unique opportunities to further define synergy and the contribution of discrete, engineered CAR-γδ T cell subsets to therapeutic efficacy *in vivo*.

## INTRODUCTION

Gamma delta (γδ) T cells are a distinct subset of T cells characterized by their expression of heterodimeric pairs of gamma and delta T cell receptor (TCR) chains^1^. Unlike alpha beta (αβ) T cells, γδ TCRs are not restricted to antigen presented on major histocompatibility complexes (MHCs) and therefore can directly bind unprocessed antigen including small molecules and proteins associated with malignant transformation and pathogenic infection^2–5^. γδ T cells can also be activated via TCR-independent mechanisms^6^ including natural killer group 2D (NKG2D), a receptor shared by NK, γδ, and αβ CD8+ T cells that binds to MHC class I polypeptide-related sequence-A and -B (MICA, MICB) and UL16-binding proteins (ULBP1)^7^, all of which display increased expression by transformed cells^8,9^. Upon activation, γδ T cells mediate their cytotoxic effects through the release of cytokines, chemokines, and cytolytic molecules, as well as upregulation of ‘death receptors’, such as Fas ligand and TRAIL^10–17^.

The importance of γδ T cells as a prognostic marker for cancer has been known for some time, with the presence of tumor-infiltrating γδ T cells identified as a favorable clinical feature for patients with colorectal cancer, head and neck squamous cell carcinoma, and gastric cancer^18–20^. Indeed, a meta-analysis of over 18,000 tumor samples from 39 different types of cancer identified γδ T cell infiltration as the most favorable immune cell predictor of improved outcomes^21^. However, clinical application of γδ T cells, and in particular γδ CAR-T cells, for adoptive cell therapy (ACT) has lagged behind αβ T cells and NK cells due to our limited understanding of their biology and a lack of effective methods for genetic modification and scalable *ex vivo* expansion. In the context of ACT, γδ T cells offer several advantages over αβ T cells or natural killer (NK) cells, including their ability to recognize a broad range of MHC-independent antigens, and the nearly absent risk of graft versus host disease (GvHD)^22–25^. Additionally, γδ T cells are widely distributed in tissues, particularly the Vδ1 subset, which exhibits superior migratory potential and function in hypoxic environments^26^. The ability of γδ T cells to function as professional antigen presenting cells (pAPCs) is often overlooked, particularly their ability to prime CD4+ and CD8+ αβ T cells and amplify adaptive immune responses^27^.

There are three prominent Vδ alleles (i.e. *TRDV1*, *TRDV2*, and *TRDV3*), which code for Vδ1, Vδ2, and Vδ3, respectively^28^. The Vδ2 subset predominates in peripheral blood and is the most studied subset, comprising 1-5% of all peripheral blood T cells.^29,30^ Of these, 50-90% are paired with the Vγ9 allele (Vγ9Vδ2 T cells)^31^. Vδ1 T cells predominate within tissues including mucosa, tongue, intestine, lung, liver, and skin and to a variable extent, the peripheral blood; particularly in patients who have been exposed to cytomegalovirus (CMV) or Epstein Barr virus (EBV)^32–35^. Vδ1-Vδ2-CD3+ T cells are a minor group that are found in blood, liver, and gut and have been shown to induce dendritic cell and B cell maturation^36^.

Zoledronate-mediated expansion of Vγ9Vδ2 γδ T cells can produce large numbers of cells that are effective *in vitro* against a variety of cancer cell lines^37–40^. However, expansion with zoledronate may upregulate exhaustion markers CD57, PD1, and TIM3 with a consequent decline in expansion after day 21^37,41^. Artificial antigen presenting cells have also been applied to expand polyclonal γδ T cells^42^; however, this approach is limited by batch-to-batch inconsistency, a 40+ day production timeline, and regulatory hurdles including the need to establish master feeder cell banks^43^. In this study, infusion of various non-engineered γδ T cell subsets in mice with ovarian cancer xenografts identified a hierarchy in terms of survival with Vδ1^+^>Vδ1^-^Vδ2^-^>Vδ2^+^, highlighting an opportunity to study the subset specific influence on survival in the context of ACT. Isolation and outgrowth of specific subsets of γδ T cells has been the subject of much recent research, with groups using plate bound anti-Vδ1 antibody to selectively stimulate and outgrow the Vδ1 subset^44^, or by using a combination of immunomagnetic separation and stimulation with anti-γδ TCR plate bound antibodies^45^. Additionally, a GMP-compliant, feeder-free protocol using cytokines and anti-CD3 antibodies selectively outgrew Vδ1 T cells with *in vitro* and *in vivo* anti-tumor cytotoxicity and robust production of IFNγ and TNFα^43^.

Genetic modification of γδ T cells through the addition of TCRs or chimeric antigen receptors (CAR) improves the cytotoxic function of γδ T cells against cancer cells^46–48^. Various iterations of γδ CAR-T cells have demonstrated 2-3 fold greater cell lysis vs. unmodified γδ T cells^49^, improved on-target-on-tumor effect^50^, improved migration to tumors^27^, and vigorous response to cancer cell lines *in vitro* and *in vivo*, including tumor rechallenge models^51^. The use of gene editing approaches like CRISPR/Cas9 to knockout (KO) negative regulators of γδ T cell function (i.e. *PDCD1*, *CISH*, *FAS*) is a powerful approach to enhance T cell function^52–54^, but is largely unexplored in γδ T cells. While the development of γδ CAR-T cells and their use in clinical trials lags behind the use of αβ T cells, it remains an active avenue of research with several clinical studies currently underway to test their efficacy and safety^55^. Another method to engineer γδ T cells with enhanced resilience is to create γδ T cells that express a dominant negative FAS receptor (FAS^DN^), as a means of evading Fas/Fas ligand signaling^56^. T cells engineered to express FAS^DN^ display increased persistence and cytotoxicity following adoptive cell transfer^56^, representing an attractive target in engineering in γδ T cells. A complementary approach to improve γδ CAR T cell persistence is to armor them using constructs that constitutively produce soluble IL-15, which promotes homeostatic immune cell proliferation and survival *in vivo*^44^. This strategy has improved antitumor activity of αβCAR T cells specific for CD19, GPC-3, CLL-1, GD2, and IL-13Ra2^57–61^. Cytokine armoring has become highly prominent in the use of NK and CAR-NK therapies, as NK cells do not persist in preclinical animal models or clinical trials without cytokine support^62,63^. Although it is not yet definitely understood if γδ T cells require cytokine support for *in vivo* persistence, particularly in the allogeneic setting, recent publications indicate that cytokine armoring improves persistence in immunodeficient animal models^51^.

Here, we present a novel, non-viral, feeder free system for the expansion and manufacturing of multiplex-edited, polyclonal CAR γδ T cells for allogeneic cancer immunotherapy. Our novel manufacturing protocol involves two rounds of stimulation and ∼22 days of expansion, achieving 15,000-fold expansion while maintaining subset polyclonality. Transposon-based integration of CD19-CAR and multiplexed base editing (BE) demonstrates up to 50% transposon integration, 5,000 fold expansion, and 99% multiplex BE. γδ CAR T cells generated using this platform are highly functional *in vitro* and *in vivo*. Thus, our current good manufacturing practices (cGMP)-compatible platform allows for high efficacy, non-viral integration of CAR and multiplex BE while retaining γδ T cell subset diversity, reducing exhaustion, and increasing tumor-specific killing.

## MATERIALS AND METHODS

### Donor γδ T Cell Isolation and Expansion

Human peripheral blood mononuclear cells (PBMCs) were obtained from healthy, de-identified donors by automated leukapheresis (Memorial Blood Centers, Minneapolis, MN, USA), then further purified via Ficoll-Hypaque (Lonza) separation. γδ TCR^+^ cells were enriched by immunomagnetic negative selection using an EasySep Human Gamma/Delta T Cell Isolation Kit (STEMCELL Technologies). After purification, γδ T cells were stimulated with either 5 μM zoledronic acid (Sigma-Aldrich) or 10 μg/mL plate-bound αPan-γδ TCR antibody (Miltenyi Biotec) and 2 μg/mL soluble αCD28 antibody (ThermoFisher). Ab-expanded populations were restimulated after 11 days using equivalent concentrations of each reagent. Plate-bound αVδ1 and αVδ2 Abs (Miltenyi) were substituted for Pan-γδ TCR when indicated to selectively outgrow specific subsets.

### Cell Culture

γδ T cells were maintained in OpTimizer T cell expansion media (Gibco) containing 10% human AB serum (Valley Biomedical), 2 mM L-glutamine (Gibco), and 100 U/mL Penicillin-Streptomycin (Millipore) and supplemented with 1,000 U/mL IL-2 (Peprotech), 5 ng/mL IL-7 (Peprotech), and 5 ng/mL IL-15 (Peprotech), referred to as “complete media” unless otherwise noted. Raji cells expressing luciferase (Raji-Luc) cells, were thawed in RPMI 1640 (Cytiva) supplemented with 10% fetal bovine serum (R&D Systems) and 100 U/mL Penicillin-Streptomycin and then conditioned for growth in OpTimizer T cell expansion media (Gibco) containing 10% human AB serum (Valley Biomedical), 2 mM L-glutamine (Gibco), and 100 U/mL Penicillin-Streptomycin (Millipore). Cell lines were validated by the University of Arizona Genetics Core using STR profiling, and were routinely tested for mycoplasma contamination (Lonza). All cells were cryopreserved at 1 × 10^7^ cells/mL in CryoStor CS10 (STEMCELL Technologies). All cell counts were performed using a Countess 3 (ThermoFisher), with trypan blue (Gibco) staining used to assess viability.

### gRNA Design

Base editor guide RNAs targeting *CISH* and *PDCD1* splice donor sites were designed using the SpliceR webtool (http://z.umn.edu/spliceR). Six NGG gRNAs per target gene were evaluated in K562 cells, and the most efficient gRNAs (**Supplementary Table 1**) were purchased from Integrated DNA Technologies (IDT) with 2’-O-methyl and 3’-phosphorothioate modifications to the first three and last three nucleotides. A gRNA designed to introduce a *FAS* dominant negative mutation was also evaluated, mimicking the naturally occurring dominant negative FAS^Y232C^ mutation^64,65^.

### Electroporation of Activated γδ T cells

Unless otherwise indicated, 48 hours after stimulation, γδ T cells were pelleted and resuspended at 4.5 × 10^7^ cells/mL in T buffer (Neon Transfection Kit; Thermo Fisher Scientific). Hyperactive *Tc Buster*^TM^ mRNA (1 μg, TriLink Biotechnologies or Aldevron) and nanoplasmid DNA encoding an αCD19 CAR (1 μg, Nature Technologies) were then added to 3.6 × 10^5^ cells on ice, followed by additional T buffer to a final volume of 12 μL. Electroporations were performed with a 10 μL Neon Pipette (Thermo Fisher Scientific) using three pulses of 1400 V and 10 ms width. To promote recovery, γδ T cells were incubated for 30 min in antibiotic-free complete media at 2 × 10^6^ cell/mL and 37°C. Cells were resuspended to a final density of 1 × 10^6^ cell/mL with complete media containing 200 U/mL Penicillin-Streptomycin. T buffer-only “Pulse” conditions were used as controls for all experiments, while nanoplasmid-only conditions were used to account for transient CAR expression. To perform multiplex gene knockouts, ABE8e Base Editor mRNA (1.5 μg, TriLink) and target gRNAs (1 μg) were added in addition to the nanoplasmid and Tc buster mRNA.

### Selection of CAR+ Cells

Following nanoplasmid transfection, stably-expressing CAR+ cells were enriched through one of two methods, depending on the selectable marker associated with each construct. CAR-γδ T cells expressing an RQR8 motif (Supplementary Figure 4A-C) were isolated by magnetic separation with αCD34 beads (Miltenyi), then assessed for RQR8 expression using flow cytometry. γδ T cells engineered with mutant *DHFR*-containing CAR constructs (Supplementary Figure 4D) were selected by adding 0.12 μg/mL methotrexate (MTX) to the culture media. Selected cells were exposed to MTX between days 5 and 11, unless otherwise specified, and were assessed for GFP expression by flow cytometry.

### Antibodies and Flow Cytometry

Unconjugated αPan-γδ TCR (Clone: REA5910; Miltenyi), αVδ1 (Clone: REA173; Miltenyi), and αVδ2 antibodies (Clone: B6; BioLegend) were used to expand γδ T cells. CAR knock-in efficiency was quantified by GFP fluorescence and surface RQR8 expression. eFluor 780-conjugated fixable viability dye (eBioscience) was used for live/dead discrimination. See **Supplementary Table 2** for antibodies used in flow cytometry. Data were collected with a CytoFLEX S flow cytometer (Beckman Coulter) and all data were analyzed with FlowJo version 10.10.0 software (BD Biosciences).

### Analysis of Gene-Editing Efficiency

Following engineering and expansion, genomic DNA was extracted using a GeneJet Purification Kit (ThermoFisher). To assess BE rates, PCR amplification was performed at the *CISH*, *PDCD1*, and *FAS* loci using primers (**Supplementary Table 3**) designed in Primer3Plus version 3.3.0^66^. Following PCR purification with a QIAquick kit (QIAGEN), samples were outsourced for Sanger sequencing (Eurofins Genomics). The resulting .ab1 files were analyzed with EditR version 1.0.0 (http://baseeditr.com/)^67^.

### CISH KO cytokine reduction assays

Day 22-expanded *CISH* KO cells were thawed, resuspended in cytokine-free media, then supplemented with IL-2 or IL-15 at the indicated concentrations. Cultures were expanded for nine days, with cell count and viability recorded every three days.

### Dominant-negative FAS assays

FAS^DN^ and triple edited cells were thawed, resuspended in complete media, seeded at 1 × 10^6^ cells/well in a 24-well plate, then rested overnight. Cells were then treated with recombinant Fas ligand (300 ng/mL; R&D Systems) and αHis-tag antibody (10 μg/mL; Clone: AD1.1.10; R&D Systems), and counted after 24 hours. A second set of KO samples was stimulated for 48 hours with αPan-γδ TCR and αCD28 prior to FasL treatment before use in *in vitro* killing assays.

### Droplet Digital PCR (ddPCR)

ddPCR was used to assess the number of stably integrated CD19 CAR constructs following γδ T cell transposition. Genomic DNA was extracted using a GeneJet Purification Kit (ThermoFisher). Primers and probes were designed using PrimerQuest online tool (Integrated DNA Technologies) using settings for 2 primers + probe quantitative PCR. Each sample was run as a duplexed assay consisting of an RNAseP internal reference primer + probe set (HEX) and a WPRE primer + probe set (FAM). Primers and probes were ordered from IDT (**Supplementary Table 4**). Reactions were set up using the ddPCR Supermix for Probes (no dUTP; Bio-Rad) with 200 ng genomic DNA per assay, according to manufacturer’s instructions in technical triplicate. Droplets were generated and analyzed using the QX200 Droplet-digital PCR system (Bio-Rad). Thermocycler conditions were 95°C for 10 min, followed by 40 cycles repeat of 94°C for 30 seconds and 60°C for 1 min, ending with 98°C for 10 min and 4°C infinite hold. Frequency was visualized and quantified via the QuantaSoft version 14.0 software (Bio-Rad). Vector copy number of averaged triplicate was corrected through initial subtraction of background from non-engineered cells followed by normalization to the percentage of γδ T cells expressing RQR8.

### *In Vitro* Killing Assays

To assess the cytotoxicity of CAR-γδ T cells, viably maintained Raji-Luc cells were seeded at 5 × 10^4^ cells/well in cytokine-free γδ T cell media across a black 96-well round-bottom plate. Cryopreserved γδ T cells expanded out to day 11 or 22 were thawed, rested overnight, titrated in cytokine-free media, then cocultured at 3:1, 1:1, 1:3, or 1:9 effector:target (E:T) ratios with Raji-Luc cells. Negative controls containing only Raji-Luc cells and positive controls treated with 1% Triton X-100 were also included. After normalizing media volumes to 200 μL, cells were incubated for defined periods of 12, 24, or 48 hr, followed by addition of 20 μL D-luciferin (10% in cytokine-free γδ T cell media). Cytotoxicity was then assessed as a function of luminosity detected by a plate reader (BioRad). To perform serial killing assays, samples displaying >90% cytotoxicity at a 3:1 E:T ratio were counted, titrated, then replated with new Raji-Luc cells every 48 hours. In all cases, samples at each E:T ratio were plated in technical triplicate.

### *In Vitro* Stimulation and Functional Assays

Surface and intracellular cytokine staining (ICS) were performed following stimulation with various concentrations of plate-coated pan-γδ TCR antibody and 2 μg/mL of soluble CD28 antibody. To coat the plate with pan-γδ TCR antibody, 100 μL of 0, 0.5, 1, 2, 5, 10, 20 μg/mL of antibody solution was added in a 96-well plate, incubated overnight at 4°C and washed twice with PBS before adding γδ T cells that had previously been expanded for 22 days. γδ T cells were seeded at 1 × 10^5^ cells/well RPMI containing 10% FBS and 1× Pen/Strep and supplemented with 2μg/mL of soluble CD28 antibody. For surface staining, the cells were incubated for 24 hrs in a 37°C incubator and stained with appropriate antibodies. For ICS, the cells were incubated for 4 hrs with GolgiPlug (BD Biosciences) and monensin (eBioscience) with cells treated with an activation cocktail (Biolegend) as a positive control. After incubation, the cells were harvested, stained with eFluor 780 and appropriate surface markers to stain the γδ T cells subset, and then permeabilized and fixed with cytofix/cytoperm solution (BD Biosciences) according to the manufacturer’s instruction and BD Perm/Wash buffer (BD Biosciences) was used for all subsequent washes and incubations. To stain intracellular cytokines, eFluor 450-conjugated αINF-γ (Clone: 4S.B3; Invitrogen) and Brilliant Violet 510-conjugated αTNF-α (Clone: MAB11; BioLegend) were added and incubated for overnight at 4°C. Data was acquired on a CytoFLEX S flow cytometer and analyzed using FlowJo software version 10.10.0 (BD Biosciences).

### *In Vivo* Tumor Challenge

NSG mice were injected intravenously (IV) with 5 × 10^5^ Raji-Luc cells. Four days later, control and engineered γδ T cells were thawed, normalized to 50% CAR positivity by addition of pulse or pulse BE-edited cells, then injected via tail vein at 10 × 10^6^ cells/mouse. Select groups received a second infusion of engineered γδ T cells seven days after the first infusion as indicated. Peripheral blood was collected weekly through the submandibular vein and tumor size was assessed via IVIS tumor imaging (Revvity). Peripheral blood, bone marrow, and other tissues (see histology) were collected at endpoint from the longest surviving mice. Blood samples were analyzed by flow cytometry and an IL-15 ELISA kit (abcam) was used to measure peripheral blood IL-15 concentration. Tumor image regions of interest (ROIs) were assessed in ImageJ version 1.54i 03 and changes in tumor burden were quantified in Python 3.11.4.

### Immunohistochemistry (IHC)

All tissues were fixed with 10% formalin, paraffin embedded, sectioned, and stained with anti-CD3 (Clone: SP7, Novus Biologicals). CD3 stain was detected using 3, 3’ diaminobenzidine (DAB) in the presence of HRP and counterstained with hematoxylin. Slides were imaged using Proscia using methods previously described with slight modifications^68^.

### TCR Sequencing

Total RNA was purified from 0.5-1 million γδ T cells using the RNeasy Plus mini kit (Qiagen) according to the manufacturer’s instructions and the TCR library was prepared using iRepertoire arm-PCR platform-RNA-iR-Complete Dual Index Primer Kit - (iRepertoire). Briefly, reverse transcription of 100 ng of total RNA was performed using the iR-RT-PCR1 enzyme mix. Then the PCR product was purified using selection beads followed by secondary amplification of the product was performed using iR-PCR2 allowing for addition of Illumina adapter sequences. Finally, libraries were purified with selection beads and sequenced using Illumina MiSeq 2X150 paired-end read length (MS-102-2002) (Illumina). The TCR sequence FASTQ files were uploaded to iRepertoire bioinformatics website for analysis. Shannon entropy, a measure of TCR diversity, was calculated as described previously^69^.

### Statistical Analysis

Student’s t-tests were performed to identify significant differences between two groups and one- or two-way ANOVAs followed by Tukey’s post hoc test were used to evaluate differences between three or more samples unless otherwise indicated. The significance level was set at α = 0.05. Mean values ± standard deviation are shown. Individual data points represent independent human donors unless otherwise specified. Statistical analyses were performed in GraphPad Prism version 9.5.

## RESULTS

### Antibody-Based Expansion of Polyclonal Peripheral Blood γδ T Cells

We sought to develop a feeder-free system to expand γδ T cells that was highly amenable to genome editing techniques. Given the well demonstrated ability of the anti-CD3 antibody clone OKT3 to strongly stimulate the expansion of human αβ T cells, we speculated that an anti-γδ TCR antibody might have a similar effect on γδ T cell expansion^70^. Thus, we tested the ability of a panel of antibodies including anti-Pan-γδ TCR, -Vδ1, -Vδ2, and a combination of -Vδ1, and -Vδ2 antibodies, as well as an OKT3 and an anti-CD16 antibody as controls in the presences or absence of anti-CD28 costimulatory antibody, to stimulate the expansion of γδ T cells from PBMCs *in vitro* **(Supplementary Figure 1A&B)**. Notably, cultures treated with anti-Vδ1 and anti-Vδ2 antibodies outgrew the Vδ1 and Vδ2 subset respectively, whereas cultures treated with anti-Pan-γδ TCR outgrew both subsets and maintained the subset composition of the input population **(Supplementary Figure 1C&D)**.

Stimulation with plate-bound αPan-γδ TCR antibody and soluble αCD28 antibody yielded 220-fold expansion, significantly greater than expansion with zoledronate, which produced an average of 90-fold expansion **(Figure 1A)**. Additionally, unlike zoledronate, which relies on myeloid cells to produce reactive phosphoantigens, TCR agonist antibodies allow for subsequent restimulations without the need for accessory cells. Thus, we examined whether repeated antibody treatments could be used to enhance overall expansion. We found that restimulation on day 11 significantly enhanced outgrowth over a three week timeframe, yielding a mean overall fold expansion of >14,000 by day 22 **(Figure 1B and Supplementary Figure 1E)**.

**Figure 1.**
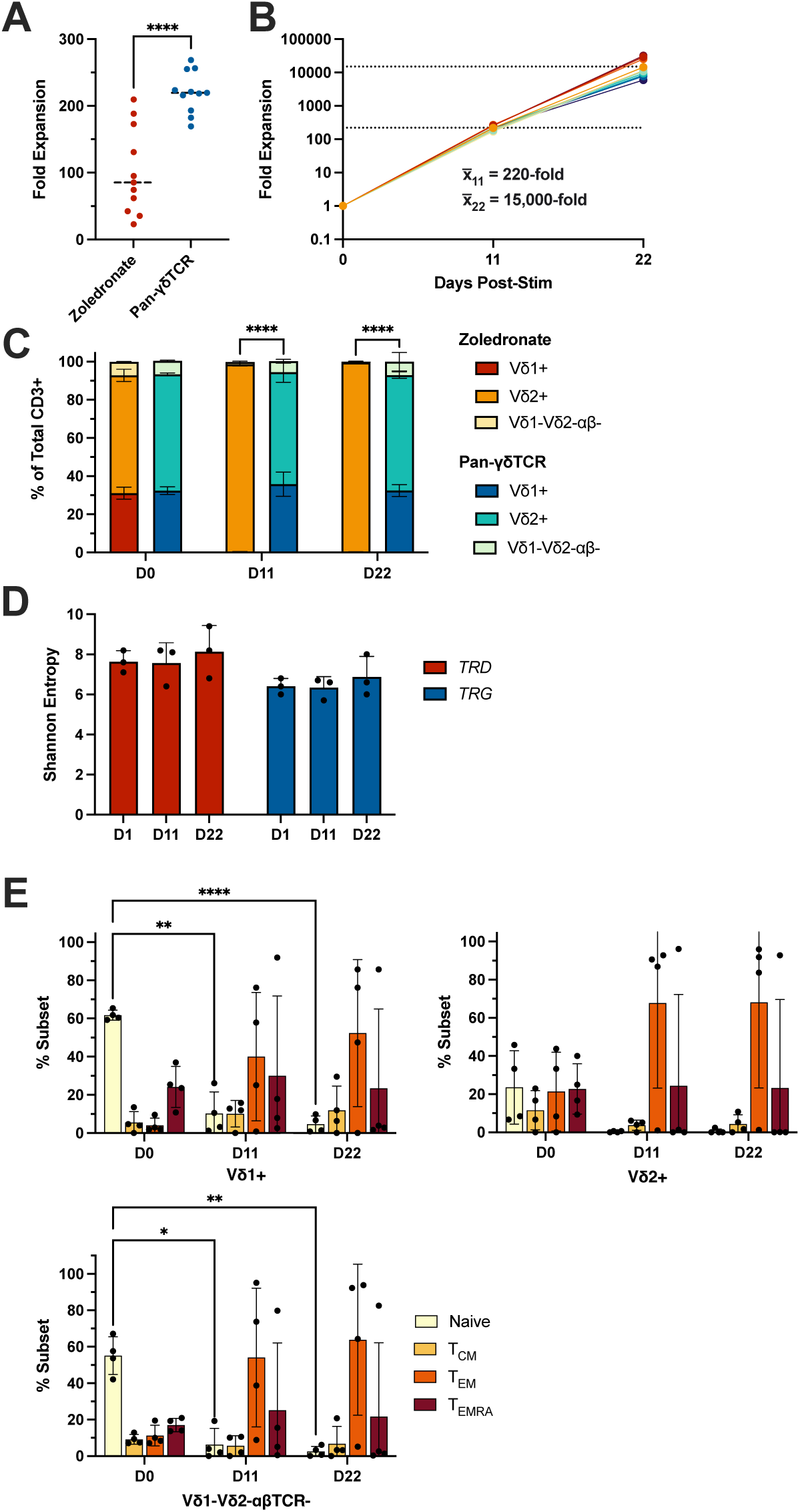
Antibody-based stimulation yields robust outgrowth of polyclonal γδ T cell populations. **(A)** Fold expansion after 11 days. Human peripheral blood γδ T cells were stimulated with either zoledronate or plate-bound αPan-γδ TCR and soluble αCD28 (n=11, paired T test). **(B)** Fold expansion at day 22 after two rounds of stimulation with αPan-γδ TCR and soluble αCD28 antibodies, occurring on day 0 and day 11 (n=11). **(C)** γδT cell subset frequency, as measured by flow cytometry, in zoledronate or αPan-γδ TCR plus soluble αCD28 stimulated γδ T cells and days 0, 11, and 22 (n=2 technical replicates of 2 independent biological donors). **(D)** Bar graphs showing Shannon entropy, a measure of TCR clonal diversity, of γδ T cell TCR diversity for TRD chains (red bars) and TRG chains (blue bars) at day 0, 11, and 22 (n=3). **(E)** Naive, T^CM^, T^EM^, and T^EMRA^ frequencies in Vδ1+ (*top left panel*), Vδ2+ (*top right panel*), and Vδ1-Vδ2-αβ- (*bottom panel*) γδ T cells subsets as determined by CD27 and CD45ra expression (n=4). (****p<0.0001, **p<0.01, and *p<0.05)

Although the functional roles of distinct γδ TCR subsets remain largely underinvestigated, current expansion protocols (zoledronate) promote outgrowth of only the Vδ2 subset^71^. Thus, we examined whether antibody-based stimulation could maintain the diversity of Vδ T cell subsets of peripheral blood γδ T cells. Unlike zoledronate-treated cells, which were almost entirely Vδ2+ by day 11, populations stimulated with αPan-γδ TCR antibody maintained consistent Vδ T cell subset frequencies across a 22 day expansion timeline **(Figure 1C)**. TCR sequencing analysis confirmed that the majority of both γ- and δ-chain diversity was maintained across each 11 day and 22 day expansion phase **(Figure 1D)**, demonstrating that antibody-based stimulation can support robust polyclonal expansion *in vitro*.

To analyze the impact of antibody-based TCR stimulation on γδ T cell maturation, we stained for naive (CD45ro+CD27+), central memory (T_CM_) (CD45ro+CD27+), effector memory (T_EM_) (CD45ro+CD27+), and effector memory re-expressing CD45ra cells (T_EMRA_) (CD45ro+CD27+) subsets^72^. The naive populations of Vδ1 and αβ TCR-Vδ1-Vδ2-subsets decreased from 61.6% ± 2.6 and 55.1% ± 10.3 on Day 0 to 10.3% ± 11.3 and 6.3% ± 8.8 on Day 11 and 4.7% ± 4.3 and 2.6% ± 2.8 on Day 22, respectively **(Figure 1E)**. Notably, our expansion protocol did not significantly impact either *TRD* or *TRG* sequence diversity or frequency (**Supplementary Figure 2**). Finally, we monitored the expression of many characteristic surface markers **(Supplementary Figure 3A**-C) over the course of our expansion protocol and found that the Vδ2 subset expressed lower levels of the exhaustion markers TIM3 and LAG3 than the either the Vδ1 or Vδ1-/Vδ2-/αβTCR-subsets and that the Vδ1 subset expressed far less NKG2A than the Vδ2- and Vδ1-/Vδ2-/αβTCR-subsets.

Taken together these data demonstrate that plate-bound αPan-γδ TCR antibody and soluble αCD28 antibody is an efficient means of rapidly expanding polyclonal γδ T cells to therapeutically relevant levels, while retaining subset and TCR diversity.

### Efficient Transposon-Mediated CAR Integration in γδ T Cells

After developing a scalable 3-week process for expanding polyclonal peripheral blood γδ T cells, we sought to enhance and direct the innate anti-tumor activity of γδ T cells by engineering them with a CD19 chimeric antigen receptor (CAR) **(Figure 2A)**. We previously developed and optimized a highly efficient non-viral CAR delivery approach in αβ T cells and NK cells using a hyperactive *Tc Buster* (TcB, BioTechne) transposon system^73^ and applied the platform to similar effect in γδ T cells **(Figure 2B)**. We found that the optimal electroporation time point for γδ T cells occurs 48 hours post stimulation, resulting in a mean stable transposition efficiency of 47.3% ± 6.1 (Mean ± SD) vs electroporating at 72 hours post-stimulation (23.7% ± 5.4) or 96 hours post-stimulation (5.1% ± 1.3) **(Figure 2C)**. Both the MTX selection **(Figure 2D)** and immunomagnetic enrichment using CD19-CAR γδ T cells transfected with a construct encoding RQR8^74^ **(Supplementary Figure 4A&B)** approaches yielded >90% CAR+ γδ T cells by day 22. Immunomagnetic enrichment did not impact production of the cytokines interferon gamma (IFNγ) or tumor necrosis factor (TNF) following stimulation with PMA and Ionomycin **(Supplementary Figure 4C).** Similarly, the frequency of CAR+ cells in non-enriched cultures did not change during expansion **(Supplementary Figure 4D)**.

**Figure 2.**
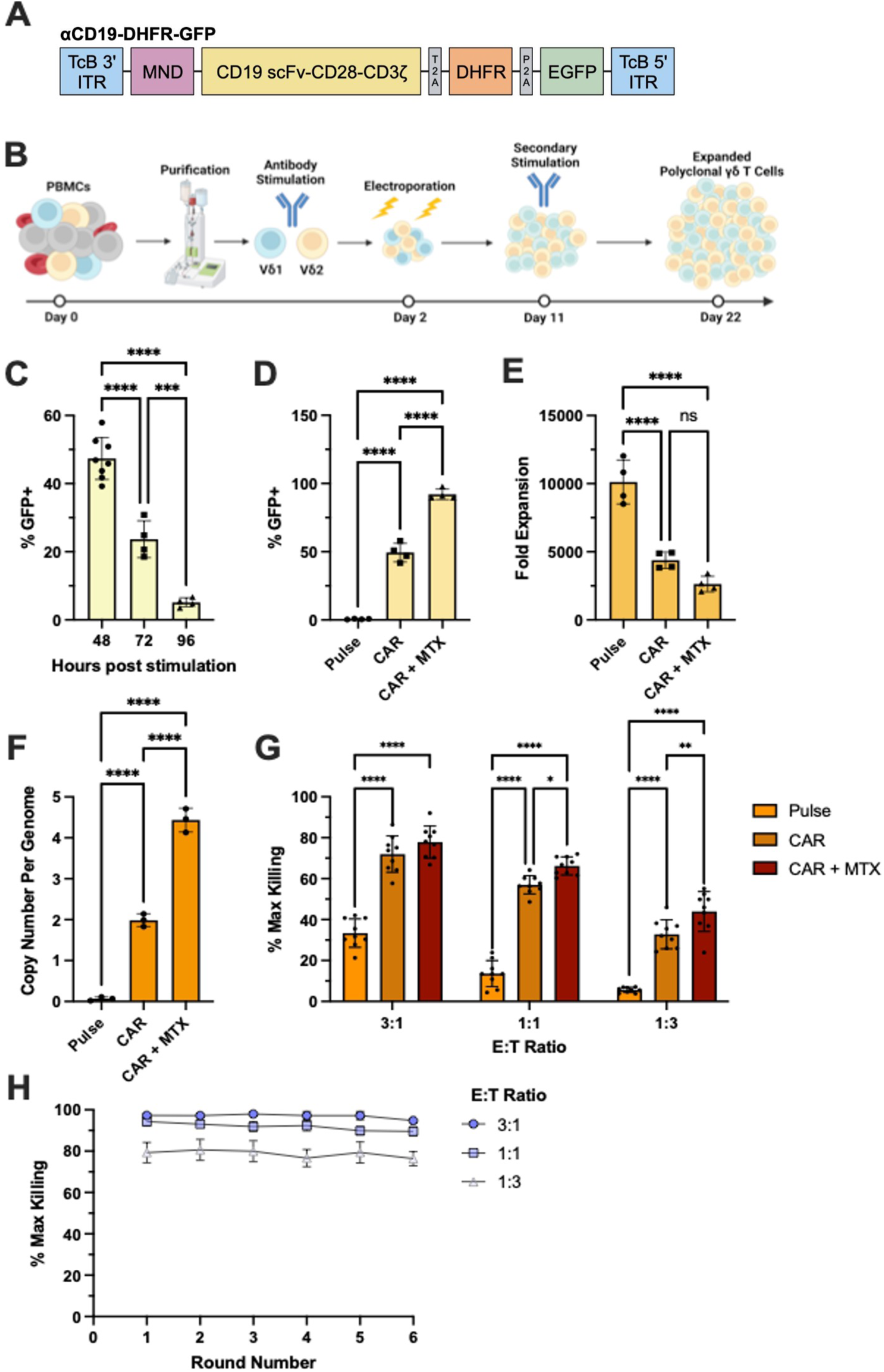
Non-virally engineered CAR-γδ T cells display potent *in vitro* cytotoxicity. **(A)** Diagram of construct containing CD19-CAR, mutant DHFR, conferring resistance to MTX in engineered cells, and GFP, used to manufacture CD19-CAR γδ T cells. **(B)** Diagram of our clinically scalable platform for γδ T cell engineering and expansion. Human γδ T cells are isolated by immunomagnetic separation from healthy donor PBMCs, then stimulated with plate-bound αPan-γδ TCR and soluble αCD28. γδ T cells are then electroporated with genome engineering reagents at day 2, re-stimulated on day 11, and collected at day 22. **(C)** Efficiency of non-viral construct delivery, as measured by GFP expression after stimulation and transfection of DNA-based CAR constructs alongside hyperactive *Tc Buster* transposase mRNA (n=8 for 48-hour time point, n=4 for 72- and 96-hour time points). **(D)** Frequency of CAR-γδ T cells, as measured by GFP expression, and **(E)** fold expansion on day 22 following MTX selection (n = 4). **(F)** CAR copy number per γδ T cell genome following non-viral integration as measured by ddPCR. Each point represents the average of three technical replicates (n=3). **(G)** *In vitro* cytolysis of Raji-luc cells after coculture with CD19-CAR γδ T cells. Assays were performed at 3:1, 1:1, and 1:3 E:T ratios in technical triplicate for two human donors. Luminescence intensity was used to quantify cytolysis of Raji-Luc target cells, with Triton X-100 serving as a positive control for max killing (n=3). **(H)** Cytotoxicity of CD19-CAR γδ T cells after repeated rounds of exposure to Raji-Luc target cells (n=6) (****p<0.0001, ***p<0.001 **p<0.01, and *p<0.05)

Notably, MTX selection or immunomagnetic selection did not impact overall viability at Day 22 **(Supplementary Figure 5A)**, nor did they impact cell health or proliferation capacity of CAR engineered cells as pulse-control cells, unselected CAR engineered cells and selected CAR engineered cells all expanded similarly during the second expansion **(Supplementary Figure 5B).**

Overall expansion in transposon engineered γδ T cells was 4,400-fold in the absence of selection and 2,600-fold in drug-selected populations **(Figure 2E)** without impacting day 22 viability or second round expansion **(Supplementary figure 5A&B)**. MTX selection also increased the average corrected vector copy number per genome from 1.99 ± 0.16 to 4.43 ± 0.29 **(Figure 2F)**, both within FDA guidelines of 5 or less transgene copies per cell^75^. Finally, CAR integration occurred at similar efficiency across Vδ1, Vδ2, and αβ TCR-Vδ1-Vδ2-subsets and the ratio of Vδ subsets were not appreciably altered by construct delivery or selection method **(Supplementary Figure 5C&D)**.

To evaluate whether CD19-CAR γδ T cells possess enhanced cytotoxicity, we performed a series of *in vitro* co-culture killing assays with target Raji-Luc cells. Because of the innate potential for non-engineered γδ T cells to exhibit CAR-independent killing, we calculated E:T ratios based on total cell number, irrespective of the percentage of CAR+ cells. While control γδ T cells largely failed to eliminate Raji-Luc cells, CD19-CAR γδ T cells completely cleared Raji-Luc cells after 24 hours at 3:1 and 1:1 E:T ratios and exhibited strong cytolysis even at 1:3 E:T ratios **(Figure 2G)**. Furthermore, D11 or D22 CAR-γδ T cells exhibited similar cytotoxicity **(Supplementary Figure 6A)**. To further assess CD19-CAR γδ T cell function we performed serial killing assays to quantify sustained cytotoxic function and monitor exhaustion, in which CD19-CAR γδ T cells that successfully cleared Raji cells were re-plated with new target cells every 48 hours successively **(Supplementary Figure 6B)**. We found that CAR-γδ T cells showed no loss of cytotoxicity throughout six rounds of killing **(Figure 2H)**, indicating robust maintenance of cytolytic function. These data demonstrate that gene engineered CAR γδ T cells can be efficiently manufactured and exhibitrobust antitumor effector function *in vitro*.

### Multiplex Base Editing of γδ T Cells

In response to stimulation, γδ T cells have been shown to upregulate FAS receptor at higher levels than αβ T cells^76^. As a result, FAS-dependent cytotoxicity has been described as a mechanism of γδ T cell activation induced cell death^77^. Provocatively, αβ T cells engineered to express FAS^DN^ transgene have shown enhanced function and persistence after adoptive transfer in solid tumor models (PMID: 30694219). Although effective, dominant negative receptor engineering requires delivery of a cDNA^78^. As an alternative, we devised an elegant strategy to utilize BE to disrupt FAS signaling by converting wildtype FAS to a FAS^DN^ (details of this will be reported separately).

To overcome activation-induced cell death in γδ T cells and the immunosuppressive tumor microenvironment we tested multiplex knock out by combining our FAS^DN^ edit with KO of *PCDC1*, a well characterized immune checkpoint, and *CISH*, a member of the suppressor of cytokine signaling (SOCS) family that provides negative feedback to gamma chain cytokine (ie IL-2, IL-4, IL-7, IL-9, IL-15, and IL-21) signaling (**Figure 3A**)^79^. Our αPan-γδ TCR antibody-based engineering protocol allowed for highly efficient BE (>90%) at all three loci, **(Figure 3B)** even when performed contemporaneously with transposon-based CAR integration **(Supplementary Figure 7A&B)**.

**Figure 3.**
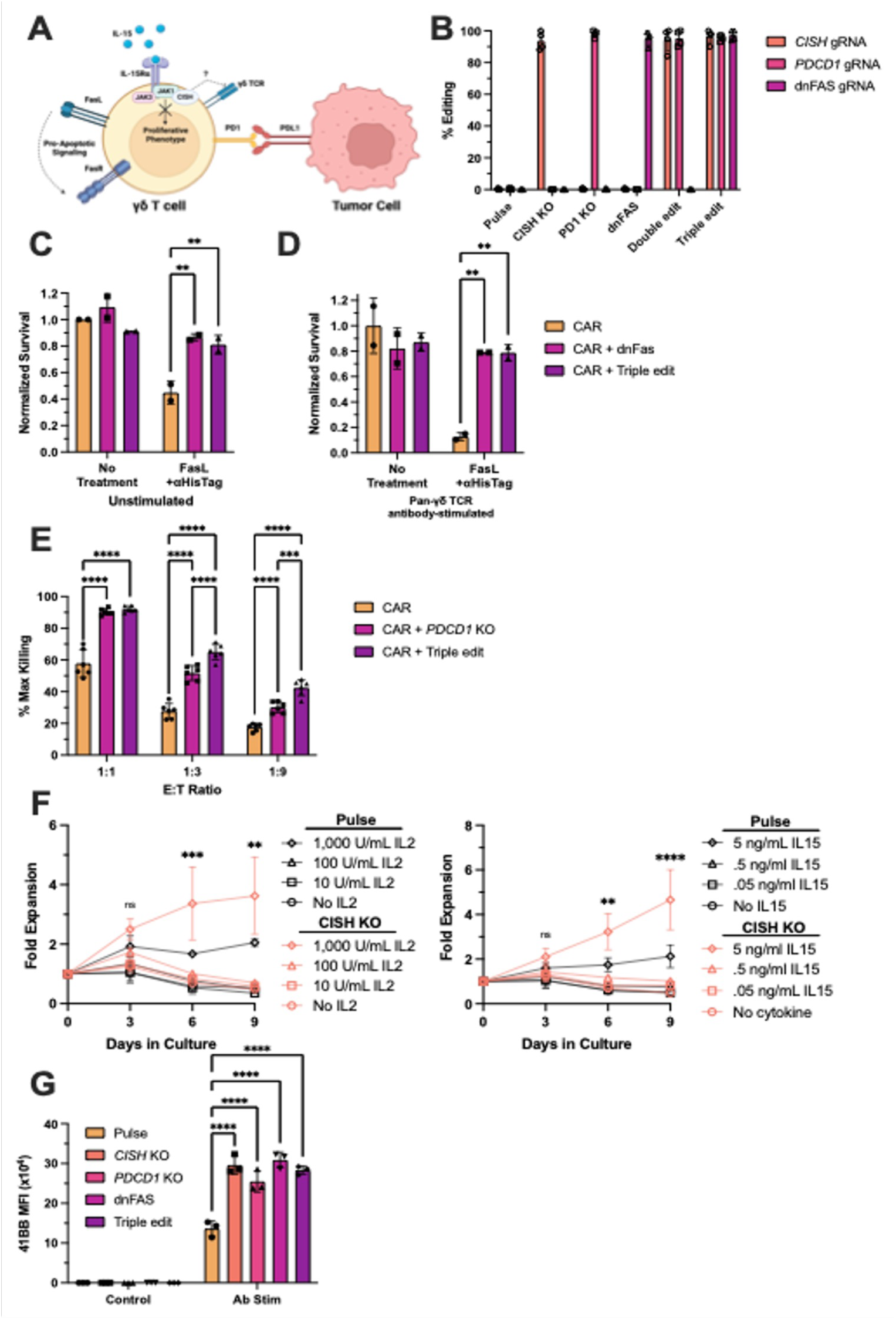
Multiplex gene knockout enhances γδ T cell function in immunosuppressive environments. **(A)** Diagram highlighting the roles of *CISH*, *PDCD1,* and *FAS* in suppressing γδ T cell activity *in vivo*. **(B)** Editing efficiency of *CISH* KO gRNA, *PDCD1* KO gRNA, dnFAS gRNA, a combination of *CISH* and *PDCD1* gRNA, and all three gRNAs, as well as a no-gRNA control in γδ T cells when cotransfected with CBE (n=4). Normalized survival of **(C)** unstimulated and **(D)** αPan-γδ TCR antibody-stimulated control, dnFAS, and triple edit CAR-γδ T cells either untreated or treated with soluble Fas ligand. Live cell counts were quantified after 24 hours and normalized to untreated control CAR γδ T cells (n=2). **(E)** Cytolysis of PDL1^high^ Raji-Luc target cells following coculture with MTX-selected CAR γδ T cells, MTX-selected CAR γδ T cells with a PD1 KO, and MTX-selected CAR γδ T cells with PD1 KO, CISH KO, and Fas dn (n=6). **(F)** Fold expansion of engineered γδ T cells cultured in in decreasing amount of IL2 (*left panel*) or IL15 (*right panel*) (n=2). **(G)** Expression of 4-1BB, as measured by flow cytometry, in gene edited γδ T cells following stimulation with αPan-γδ TCR and soluble αCD28 for 24 hours (n=3).

In the absence of TCR stimulation, γδ T cells engineered with FAS^DN^ exhibited a normalized survival of 80%, compared with 45% for CAR only controls **(Figure 3C)**. The difference was even more striking following treatment with αPan-γδ TCR, as disruption of FAS receptor signaling yielded a six-fold increase in survival **(Figure 3D)**.

To quantify the impact of PD1 KO on γδ T cells, we exposed PD1 KO and control engineered CD19-CAR engineered γδ T cells to PDL1^high^ Raji-Luc cells (**Supplementary Figure 8**) in a killing assay. As expected, CD19-CAR γδ T cells containing a PD1 KO exhibited significantly greater killing than CAR-only controls after 24 hours **(Figure 3E)**.

As previous reports found that CISH KO enhances αβ T cell responsiveness, we treated CISH KO γδ T cells with decreasing concentrations of either IL-2 or IL-15 and assessed proliferation rates, which demonstrated that expanded CISH KO γδ T cells achieved significantly greater outgrowth than untransfected pulse cells **(Figure 3F)**. This parallels the effects of loss of CISH signaling in αβ T cells and NK cells, where it acts as an important suppressor of cytokine signaling^80–82^.

Finally, we combined all three base edits to assess their cumulative effect on CD19-CAR γδ T cell phenotype and function. Triple edited CAR-γδ T cells performed equally as well as FAS^DN^-only CAR-γδ T cell in a FasL apoptosis assay **(Figure 3C&D)** and better than CAR-γδ T containing only a PD1 KO in a killing assay at low E:T ratios. **(Figure 3E)**. Following αPan-γδ TCR stimulation, all engineered cell types experienced two- to threefold greater activation as measured by 41BB upregulation relative to controls **(Figure 3G)**. Overall, these data demonstrate that targeted BE of negative regulators of γδ T cells can increase their resistance to immunosuppressive signaling while enhancing γδ T cell activation.

### Engineered γδ T Cells Display Potent *In Vivo* Cytotoxicity

To investigate the potential of expanded polyclonal γδ T cells for ACT, we utilized a well established Raji-Luc murine xenograft model^54,83^. Here, we tested groups with or without triple BE edits and with or without cytokine armoring with IL-15 expressing CD19-CAR transposons **(Figure 4A, Supplementary Figure 9A)**. We also included two extra groups that received two doses of the CD19-CAR γδ T cells seven days apart. Immunodeficient NSG (NOD.Cg-Prkdcscid Il2rgtm1Wjl/SzJ) mice were intravenously engrafted with 5 × 10^5^ Raji-Luc cells four days prior to intravenous infusion using 10 × 10^6^ expanded γδ T cells manufactured in five technical replicates from a single human donor **(Supplementary Figure 9B**-F).

**Figure 4.**
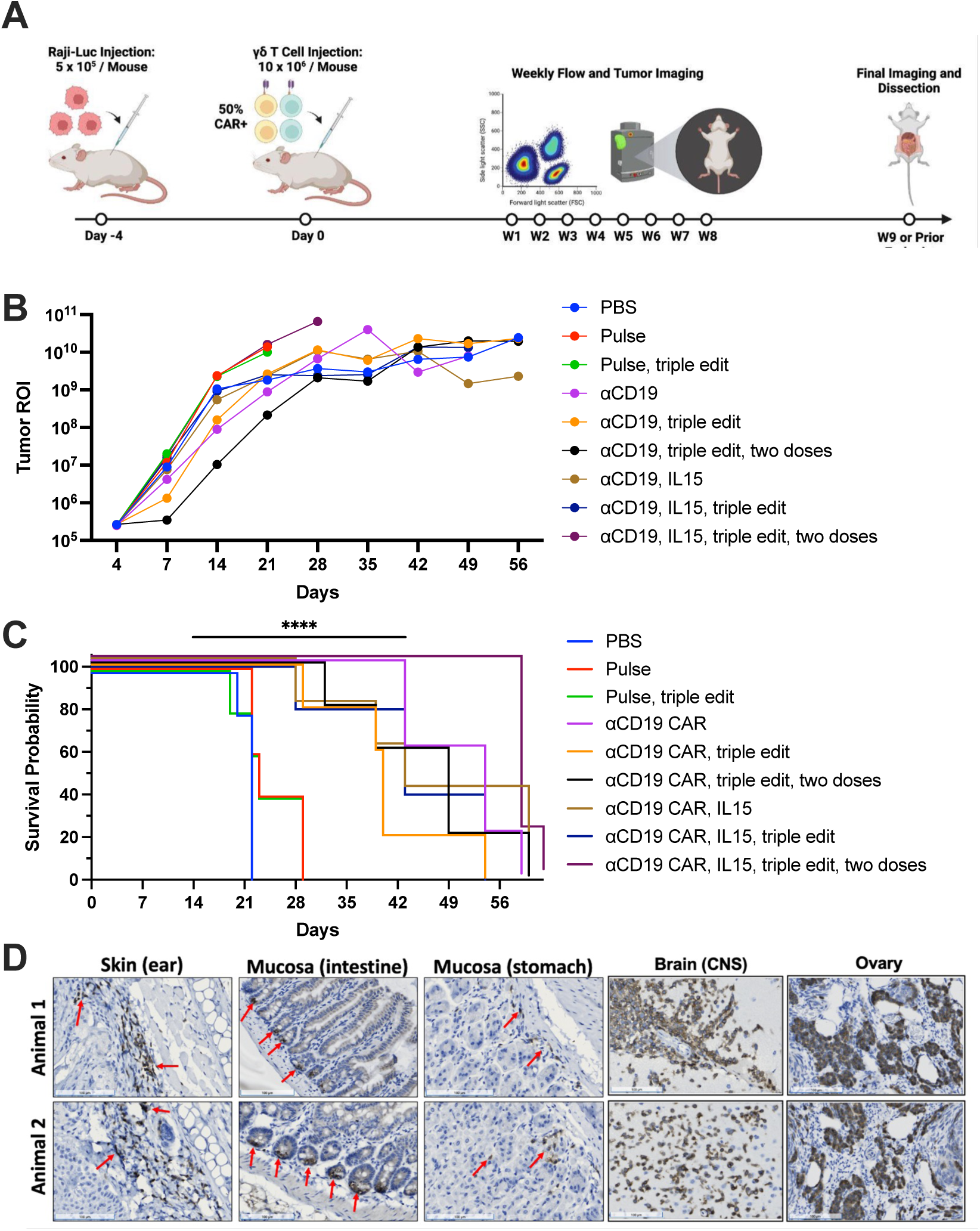
Engineered polyclonal γδ T cells exhibit potent anti-tumor activity *in vivo*. (A) Diagram of *in vivo* tumor challenge model. Mice were injected intravenously with 5×10^6^ Raji-luc cells on day -4 and treated with 10×10^6^ γδ T cells on day 0. Mice that received two doses were injected again with 10×10^6^ γδ T cells on day seven. Tumor growth and overall survival was measured and flow cytometry on blood was done weekly until they reached endpoint. (B) Tumor growth and (C) Kaplan-Meier survival curve for NSG mice injected with Raji-Luc cells and treated with the indicated γδ T cells (n=5). (D) Immunohistochemistry staining of CD3 on paraffin embedded skin, intestines, stomach, brain, and ovary in two animals treated with γδ T cells engineered with CD19-CAR and IL15.

We observed tumor growth via bioluminescent imaging weekly for the duration of the experiment. Mice treated with CAR-γδ T cells experienced suppressed tumor growth and lived twice as long as their control counterparts **(Figure 4 B&C, Supplementary Figure 10 A&B)**. Notably, mice that received engineered γδ T cells possessing an anti-CD19 CAR construct that also encoded IL-15 and triple BE edits survived nearly three times as long as mice treated with wildtype γδ T cells.

To examine the distribution of engineered γδ T cells within the treated mice, we performed necropies as mice reached the predefined endpoint. We observed robust γδ T cell infiltration, as measured by immunohistochemistry staining of human CD3 positivity in the skin, gut mucosa, and ovaries, as well as other tissues (**Figure 4D, Supplementary Figure 11**). Importantly, we also identified extensive γδ T cell infiltration in the CNS, suggesting that these cells can cross the blood-brain barrier and may have potential as a therapeutic approach for tumors in immune privileged sites (**Figure 4D**).

Overall, our data demonstrate that γδ T cells can be non-virally engineered to express a CAR with simultaneous multiplex BE without sacrificing performance or function following our cGMP-compatible αPan-γδ TCR antibody-based engineering process.

### Dynamics of Engineered γδ T Cell Subsets *In Vivo*

Following *in vitro* engineering and expansion, the ACT infusion products used for *in vivo* studies exhibited a typical peripheral blood γδ T cell subset distribution characterized by >80% Vδ2 expression across all conditions **(Supplementary Figure 8E).** As our expansion protocol does not alter the frequency of γδ T cell subsets, we monitored the dynamics of each γδ T cell subset across the *in vivo* study via weekly blood draws and flow cytometry analysis. We examined the relative fraction of each major γδ T cell subset within human CD3+ cells **(Figure 5A, Supplementary Figure 12)**, as well as their overall frequency within peripheral blood **(Figure 5B, Supplementary Figure 13A)** and bone marrow **(Supplementary Figure 13B)**.

**Figure 5.**
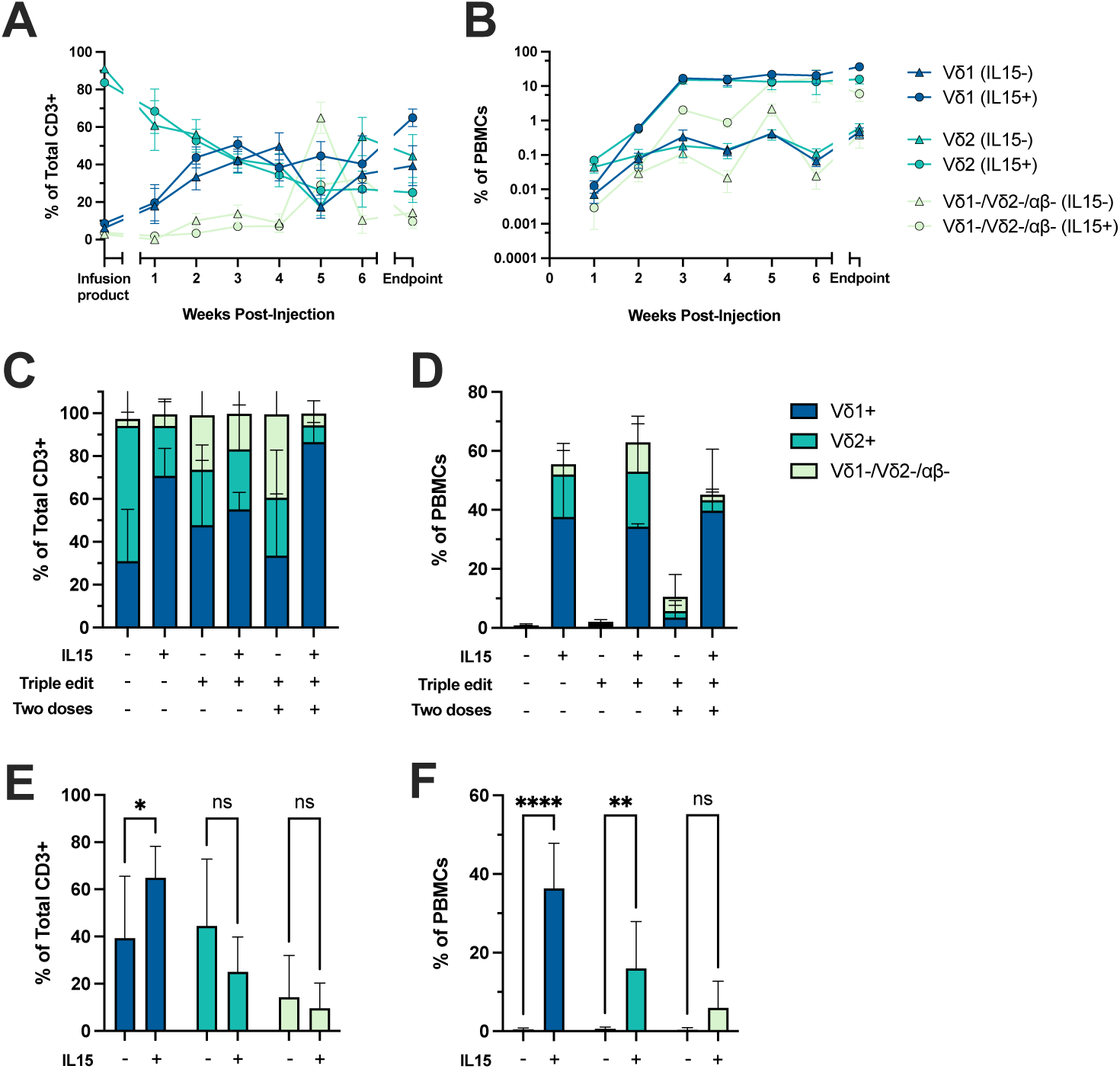
Effect of gene engineering on γδ T cell subset frequency and persistence *in vivo*. Weekly frequency of Vδ1+, Vδ2+, and Vδ1-Vδ2-αβ- subsets **(A)** as a percent of total CD3+ cells and **(B)** as a percent of total PBMC as measured by flow cytometry. “Infusion product” indicates the frequency before injection while “Endpoint” indicates the frequency at necropsy (n=10). **(C)** Frequency of γδ T cell subsets as a percent of total CD3+ cells and **(D)** as a percent of total PBMCs at necropsy as measured by flow cytometry (n=5). **(E)** Frequency of γδ T cell subsets as a percent of total CD3+ cells and **(F)** as a percent of total PBMCs at endpoint in groups that received a single dose of cells engineered with a constructs that did not contain IL15 compared with groups that received a single dose of cells engineered with constructs that contained IL15 (n=10).

We found a striking change in the relative frequency of distinct γδ T cell subsets as a fraction of total CD3+ cells, changing from ∼90% Vδ2+ and ∼10% Vδ1+ at injection to ∼50% Vδ2+ and ∼40% Vδ1+ by week two, a trend that continued throughout the study **(Figure 5)**. This increase in Vδ1+ cells and decrease in Vδ2+ cells persisted at the endpoint in the majority of challenged mice, including those that received γδ T cells without CAR engineering (**Figure 5C, Supplementary Figure 14)**.

We also observed a major difference in the overall frequency of γδ T cell subsets in peripheral blood between groups receiving γδ T cells engineered with constructs encoding IL-15 increasing to over 10% of the overall PBMCs and those that did not express IL-15 remaining below 1% **(Figure 5B&D)**. This increase occurred in all subsets with the Vδ1+ subset increasing ∼80-fold and the Vδ2+ subset increasing ∼30-fold between the first week post-infusion and endpoint (**Supplementary Figure 15**). However, despite this notable difference, there was no improvement in survival in groups that received IL-15 secreting cells versus those receiving CAR-engineered cells alone **(Supplementary Figure 16)**.

Additionally we performed TCR sequencing on the infusion product and two of the mice belonging to the αCD19-CAR triple BE edit w/ IL15 cohort at endpoint. Notably, the frequency of the top 10 TCR clones for both the *hTRG* and *hTRD* genes present in the infusion product are greatly reduced or absent from T cells in the blood and bone marrow **(Supplementary Figure 17A&B, *left column*)**. Similarly, the frequency of the top 10 genes present in the blood and bone marrow are also absent or greatly reduced in frequency when compared with the infusion product (**Supplementary Figure 17A&B, *center and left columns)***. This coincides with an increase in overall γδ T cell subset diversity in the blood and bone marrow after infusion (**Supplementary Figure 18**).

The inclusion of IL-15 had the largest impact on the Vδ1+ population, increasing both the fraction of Vδ1+ cells as a percent of CD3+ cells **(Figure 5E)** and as a percent of total PBMCs **(Figure 5F)**. The decrease of Vδ2+ as a frequency of CD3+ in the presence of IL-15 did not reach significance, but the increase of Vδ2+ as a frequency of total PBMCs did.

Notably, the frequency of γδ T cells expressing the CD19-CAR, as measured by RQR8 expression, increased throughout the experiment in mice treated with cells engineered with a CD19-CAR, cells engineered with a CD19-CAR and a triple BE edit, and cells engineered with a CD19-CAR and IL-15 **(Supplementary Figure 19)**, indicating that CAR+ cells may have an expansion or survival advantage in an *in vivo* tumor challenge model. However, mice treated with cells engineered with a CD19-CAR, a triple BE edit, and IL-15 showed no significant change in frequency of cells expressing CD19-CAR. This lack of change could be attributed to a lack of statistical power or ineffective synergy between the combination of genetic changes.

Taken together, these data demonstrate that CAR γδ T cells generated with αPan-γδ TCR antibody-based engineering are functionally active *in vivo* and can be used to help control tumor growth. It also shows that IL-15 can greatly increase the baseline level of both Vδ1+ and Vδ2+ γδ T cells in peripheral blood, with greater expansion of the Vδ1+ subset.

## DISCUSSION

The role of γδ T cells in cancer immunity has been the subject of intense research recently. Their ability to rapidly activate, produce proinflammatory cytokines, directly lyse target cells, and present antigen afford them a unique combination of anti-tumor capabilities. γδ T cells can be activated directly through their TCRs, independent of MHC presentation or through receptors such as TLRs, Fc receptors, or other innate immune receptors. Their activating ligands can be peptides, but also include phosphoantigens and stress related antigens such as heat shock proteins and MICA/MICB. Notably, the presence of γδ T cells in the tumor is the only cell subset that correlates with reduced disease progression^19,21^.

γδ T cells possess many attractive features as a platform for ACT, including their lack of MHC restriction, potent cytotoxicity, and ability to act as professional antigen presenting cells (pAPC). Despite this, their development as a clinical option for the treatment of cancer has greatly lagged behind αβ T cells and NK cells, in part a consequence of immature methods for isolation, engineering, and expansion. This lack of development is partially due to a lack of functional knowledge and familiarity, as γδ T cells are far less well studied than αβ T cells and NK cells, and partially caused by practical issues, as γδ T cells are a relatively rare subset in the peripheral blood and murine γδ T cells are very different than human γδ T cells^84^.

One of their most compelling attributes, a unique functional diversity driven by distinct γδ TCR combinations, is attenuated by existing engineering and expansion methods^25^. Here we show an antibody-based, cGMP-compatible, *ex vivo* expansion protocol promotes the robust *ex vivo* outgrowth to clinically-relevant numbers of Vδ1, Vδ2, and Vδ1-/Vδ2-/αβTCR-cells, and is amenable to CAR integration and multiplex genome editing using Cas9 base editors. This pan-expansion of γδ T cells subsets also leads to the preservation of significant TCR diversity, enabling fundamental investigation into subset biology and inter-subset synergy.

TcBuster transposon-based integration of CAR constructs yielded roughly 50% CAR+ cells with a copy-number rate of less than five copies per genome, with good antitumor potency even at low E:T ratios. Importantly, CAR γδ T cells manufactured with this method did not show any reduction in potency over six rounds of serial killing *in vitro*, even at the lowest E:T ratios, suggesting that they are highly resistant to antigen-induced exhaustion. This is a major advantage in the setting of anti-tumor immunity as T cell exhaustion is a major contributor to loss of immune control of tumor.

Using BE, we successfully introduced precise edits at three genes (*CISH*, *PDCD1*, and *FAS*) simultaneously at rates of over 90%. PD1 KO increased the cells’ ability to lyse PDL1-expressing tumor cells while CISH KO enhanced cell persistence in low cytokine environments. Whereas KO of PD1 and CISH was accomplished via splice site disruption, we instead chose to leverage the nucleotide editing capability of base editor to introduce a previously identified dominant negative mutation at the endogenous *FAS* locus, greatly reducing the cells’ susceptibility to FAS/FAS ligand mediated suppression without the need to deliver an exogenous cDNA. We are currently adapting this approach to other suppressive surface proteins for which dominant negative mutations have been described, the results of which will be reported in greater detail separately.

The adoptive transfer of CD19-CAR γδ T cells in mice seeded with CD19-expressing Raji cells slowed tumor growth and extended survival over mice treated with PBS or non-engineered γδ T cells, indicating that CAR+ γδ T cells can contribute to tumor control. CD3+ γδ T cells were present in all tissues sampled, indicating their robust ability to traffic to and survey diverse tissues, including the CNS. Importantly, many CD3+ γδ T cells were observed in the brain, demonstrating the ability of γδ T cells to cross the blood brain barrier and highlight their potential as a therapy for both primary tumors of the CNS and for other cancers that may metastasize to the brain.

The inclusion of soluble IL-15 “armoring” in CAR constructs had a significant impact on the outgrowth of γδ T cells *in vivo*. γδ T cells that were engineered with constructs expressing soluble IL-15 exhibited substantially increased frequency in the peripheral blood, with all treatment groups containing IL-15 having over 50% of all PBMCs being γδ T cells. This increase was due to an expansion of both the Vδ1 and Vδ2 subset suggesting that IL-15 armoring may support cell engraftment and persistence in a clinical setting. However it is notable that the Vδ1 subset demonstrated greater outgrowth *in vivo* as measured by an increase in the frequency of Vδ1+ cells as a fraction of CD3+ cells and greater total fold-expansion in the peripheral blood.

γδ T cells are a group of versatile, multifunctional lymphocytes that hold great promise as an immunotherapy. Our cGMP-translatable manufacturing protocol supports further translation-focused efforts to evaluate their therapeutic potential. Further research is needed to determine the relative contribution and function of each γδ T cell subset to an overall immune response, and whether specific genetic modifications are more well-suited to enhance the function of discrete subsets. Given γδ T cells’ ability to interact and influence other immune cell types via proinflammatory cytokine production, ADCC, and antigen presentation on MHC Class I and Class II, future research using immunocompetent models is warranted to achieve the full potential of γδ T cells as a robust cellular chassis for cancer immunotherapy and beyond.

## Acknowledgments

B.R.W. acknowledges funding from Office of Discovery and Translation, NIH grants R21CA237789, R21AI163731, P01CA254849, P50CA136393, U54CA268069, R01AI146009, and, Children’s Cancer Research Fund. B.S.M. acknowledges funding from NIH grants R01AI146009, R01AI161017, P01CA254849, P50CA136393, U24OD026641, U54CA232561, P30CA077598, U54CA268069, Children’s Cancer Research Fund, the Fanconi Anemia Research Fund, and the Randy Shaver Cancer and Community Fund.

