## Supplementary Figures for "Efficient multiplex non-viral engineering and expansion of polyclonal γδ CAR-T cells for immunotherapy"

**Supplementary Figure 1**

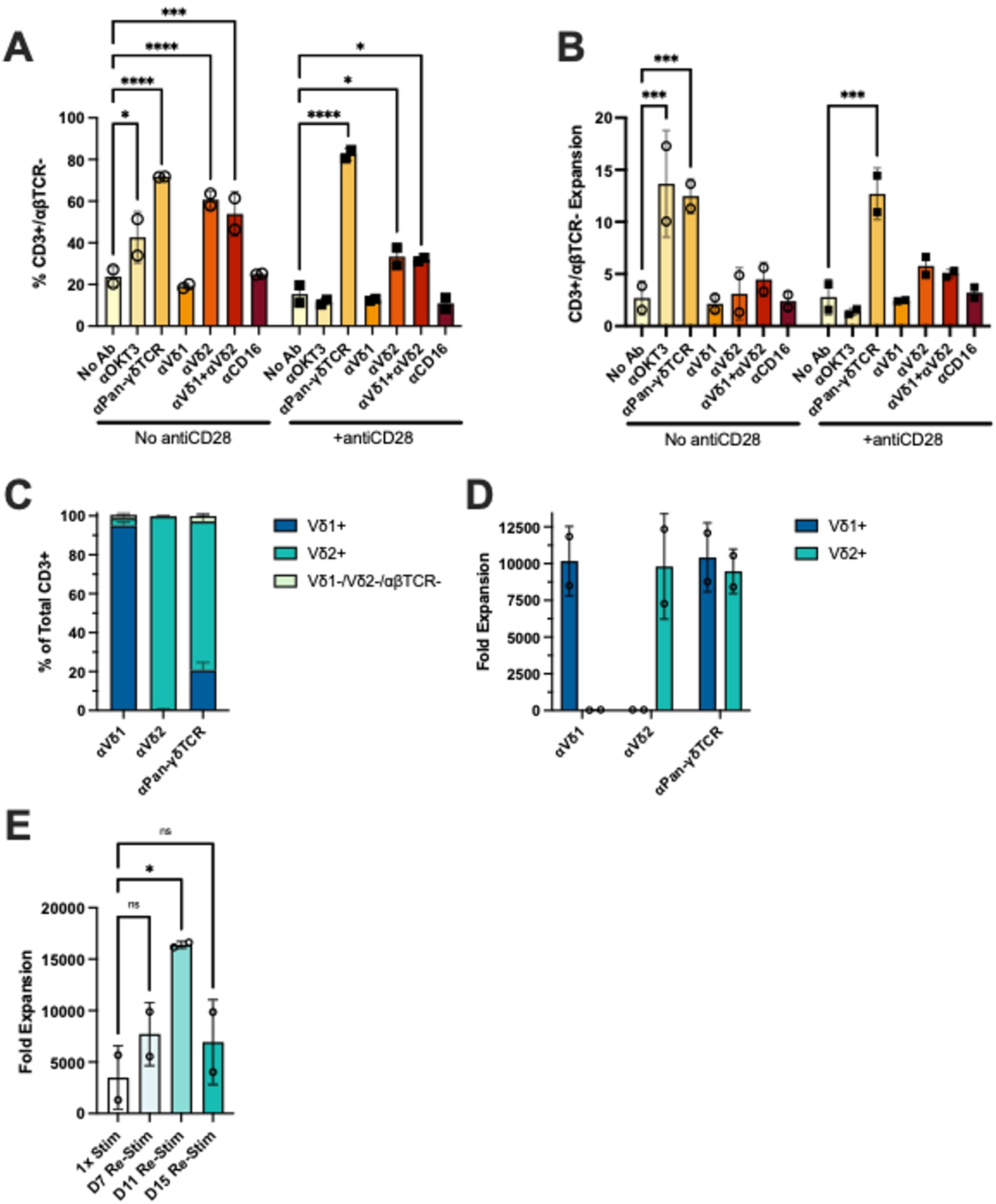


**Supplementary Figure 1. Screening of putative stimulatory antibodies identifies αPan-γδTCR as a driver of robust polyclonal γδ T cell expansion.**

**(A)** Fraction and **(B)** fold expansion of CD3+/αβTCR- cells in total PBMCs expanded for 7 days with the indicated plate-bound antibodies in the presence or absence of soluble αCD28. Significance was assessed using Student's T-test with Tukey post-hoc correction relative to the "no antibody" condition (n=2). **(C)** Relative fraction and **(D)** fold expansion of γδTCR subsets at day 22 in 2 human donors stimulated with αVδ1, αVδ2, or αPan-γδTCR, along with soluble αCD28 (n=2). **(E)** Total fold expansion of isolated γδ T cells stimulated initially with αPan-γδTCR and soluble αCD28, then re-stimulated at the indicated time points (n=2).

**Supplementary Figure 2**

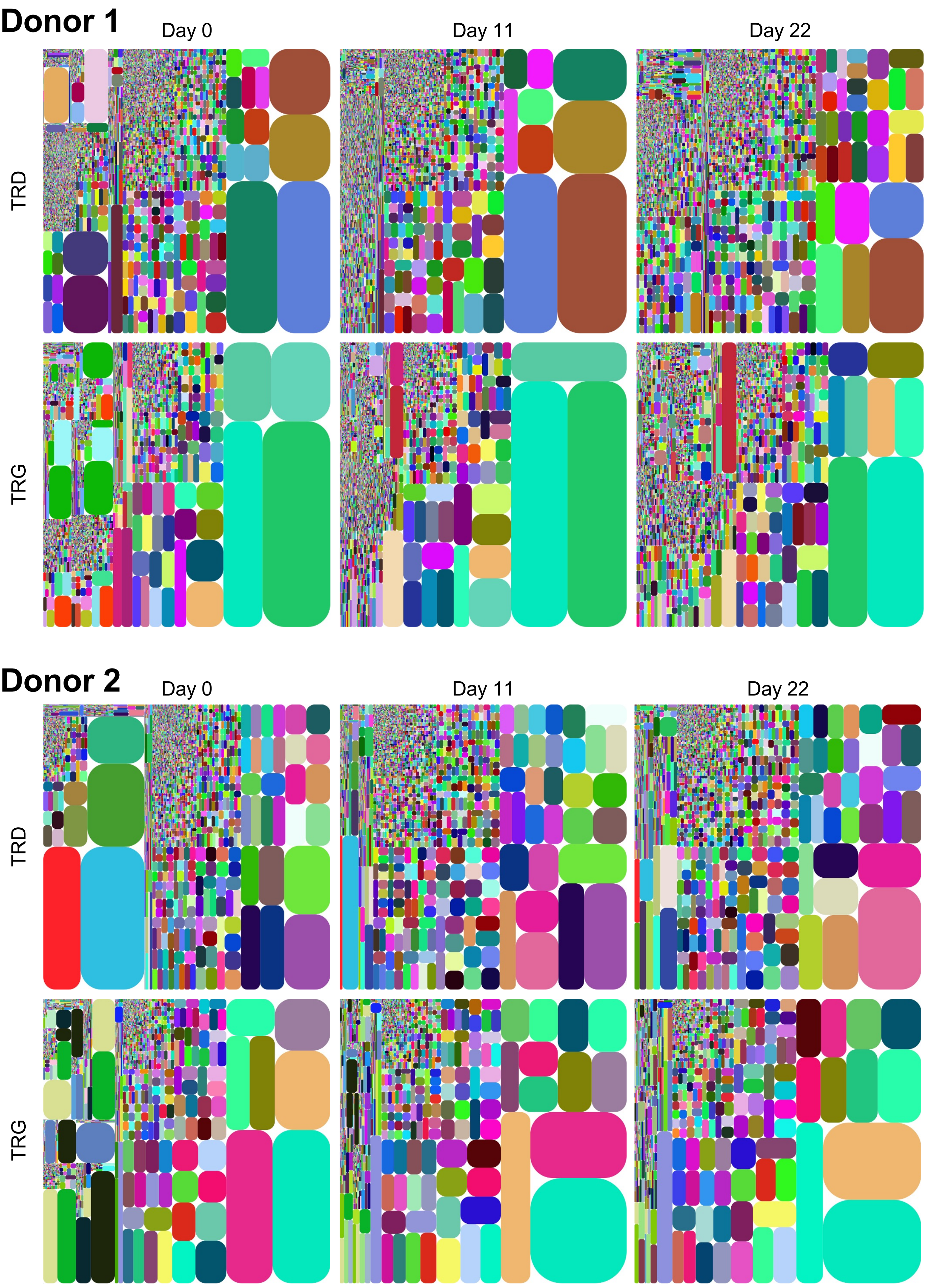

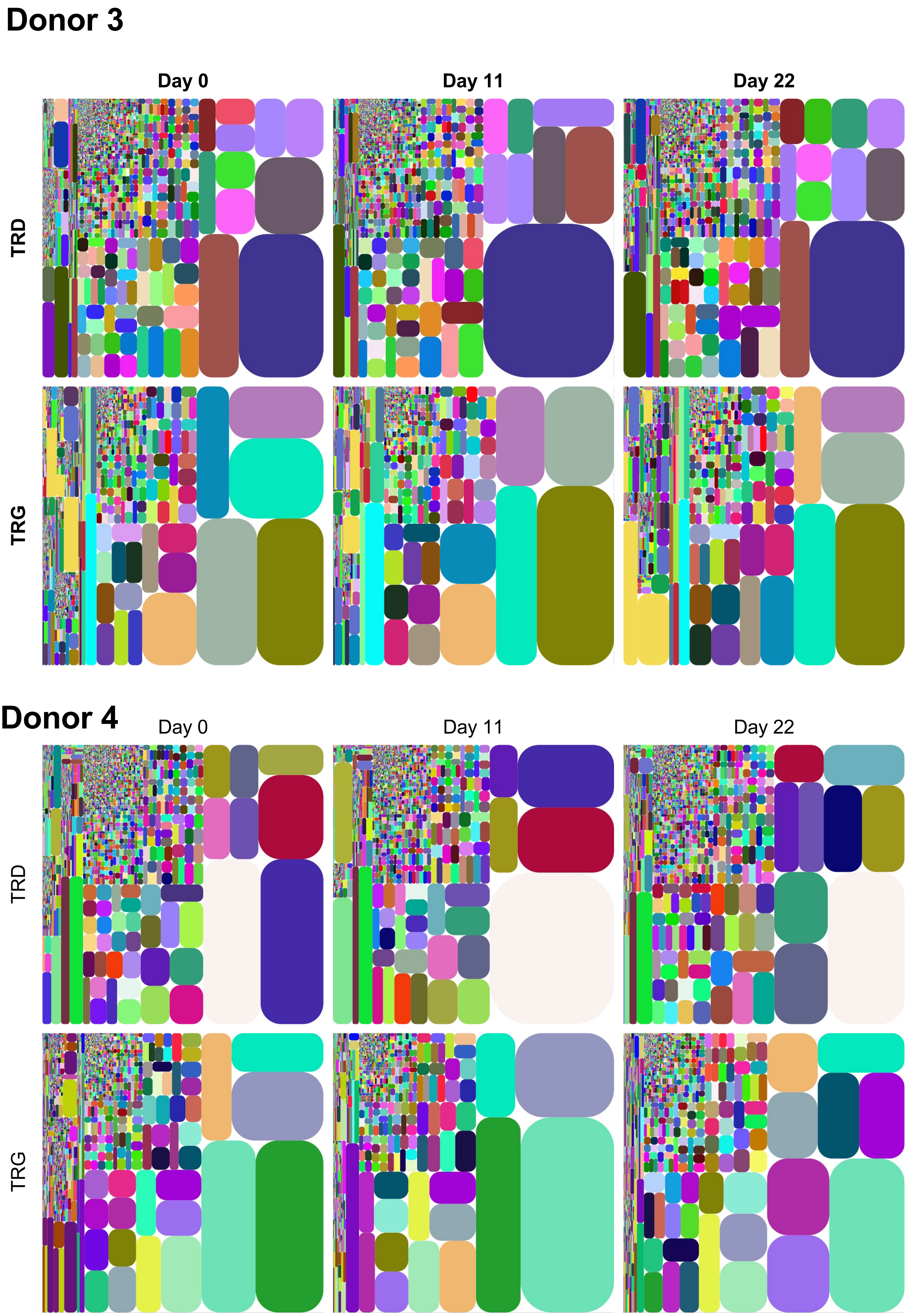
Donor 3
Day 0
Day 11
Day 22
TRD
TRG
Donor 4
Day 0
Day 11
Day 22
TRD
TRG

**Supplementary Figure 2. Treemaps of TCR *TRD* and *TRG* sequence frequency from 4 donors during αPan-γδTCR stimulated expansion**

**Supplementary Figure 3**

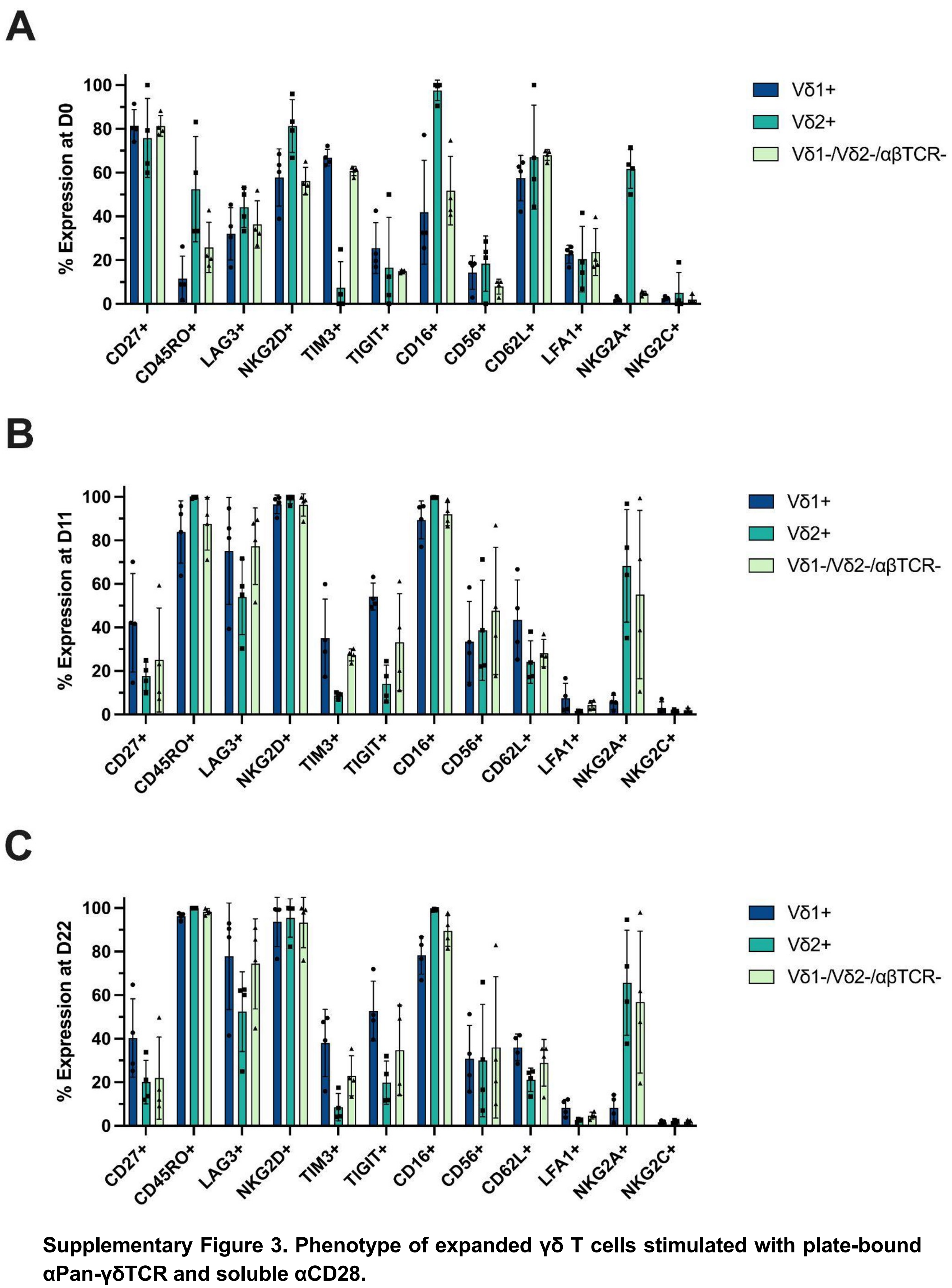


**Supplementary Figure 3. Phenotype of expanded γδ T cells stimulated with plate-bound αPan-γδTCR and soluble αCD28.**

γδ T cells were isolated by negative selection from four human PBMC donors. Surface marker expression was quantified at **(A)** day 0, **(B)** day 11, and **(C)** day 22 (n=4).

#### Supplementary Figure 4

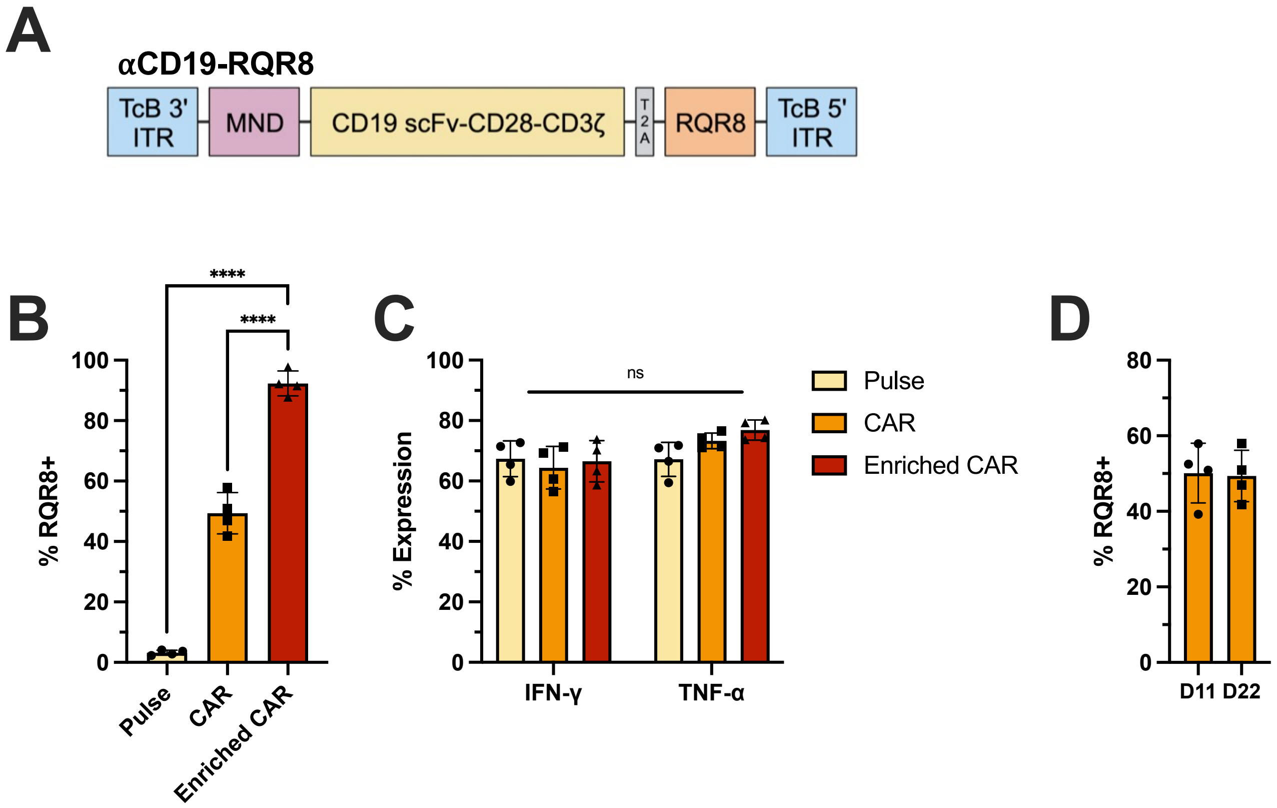


**Supplementary Figure 4. Transposon mediated integration**.
**(A)** Diagram of the construct used to manufacture αCD19-RQR8 γδ T cells. For the αCD19-RQR8 construct an MND promoter drives the expression of an αCD19-CAR, composed of an αCD19 scFv, a CD28 transmembrane domain, and a CD3ζ intracellular signaling domain and an RQR8 marker which can be used to determine the frequency of stably integrated cells or for immunomagnetic separation, flanked by flanked by 3' and 5' ITR sites, which are recognized by the *Tc Buster* transposase. **(B)** Expression of RQR8 following immunomagnetic enrichment (n=4) on αCD19-RQR8 γδ T cells. **(C)** Expression of IFNγ and TNF as measured by ICS following stimulation with PMA and Ionomycin. (n=4). **(D)** RQR8 expression at days 11 and 22 in αCD19-RQR8 γδ T cells.

### Supplementary Figure 5

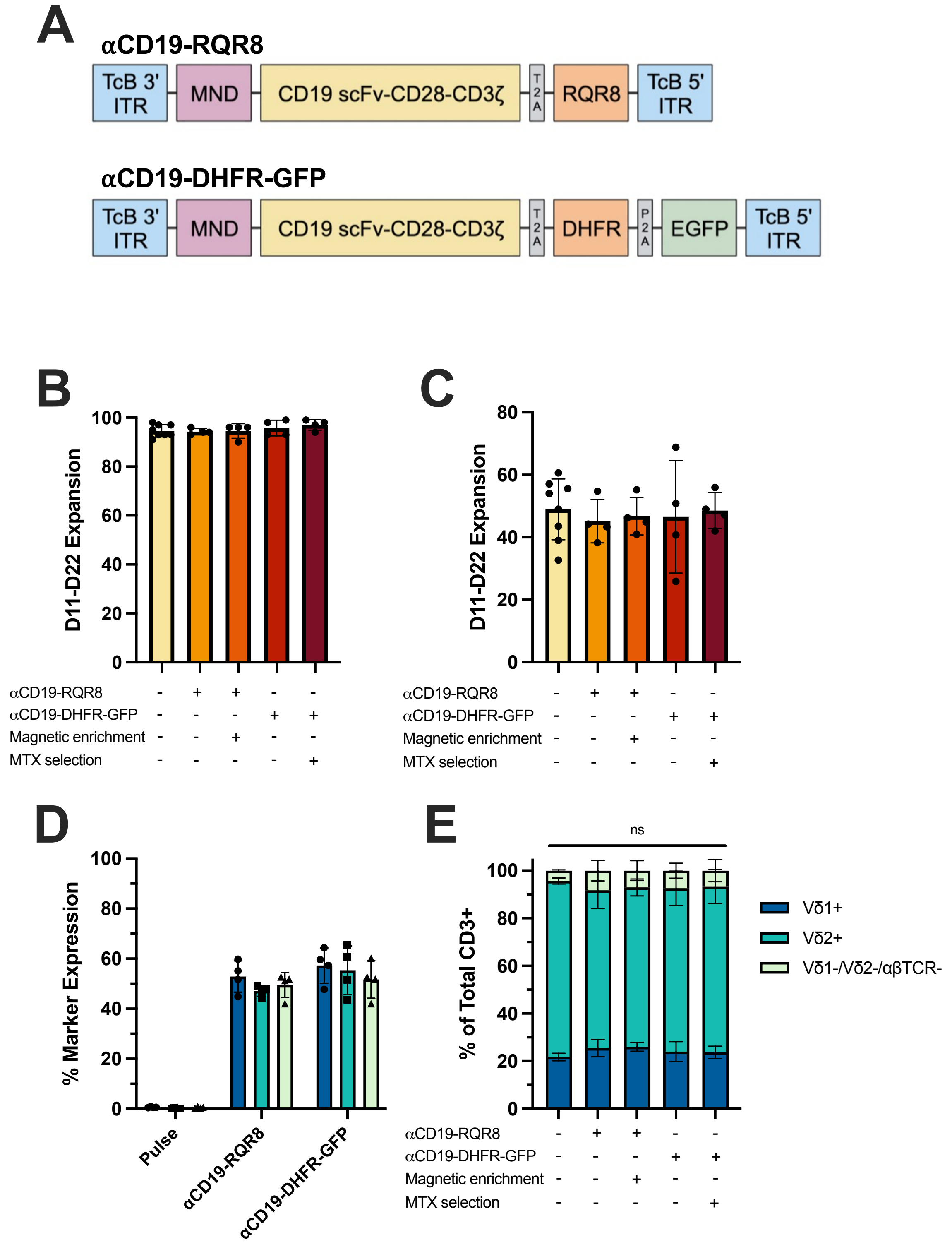

**Supplementary Figure 5. αPan-γδTCR-based expansion allows for efficient non-viral CAR integration and subsequent selection.**

**(A)** Constructs used in this figure. For the αCD19-DHFR-GFP construct an MND promoter drives the expression of a CD19-CAR, a mutant DHFR confers resistance to MTX, and GFP can be used to determine the frequency of stably integrated cells. Both constructs are flanked by 3' and 5' ITR sites, which are recognized by the *Tc Buster* transposase. **(B)** Viability at Day 22 following a second round of αPan-γδTCR-based expansion of cells transfected with αCD19 or αCD19-DHFR-GFP (n=4). The latter were selected through exposure to 0.12 µg/mL methotrexate between day 5 and day 11. **(C)** Fold expansion between days 11 and 22 following a second round of αPan-γδTCR-based expansion of cells transfected with αCD19-RQR8 or αCD19-DHFR-GFP γδ T cells (n=4). **(D)** Expression of RQR8 or GFP in Vδ1+, Vδ2+, and Vδ1+/Vδ2+/αβTCR- subsets in αCD19-RQR8 or αCD19-DHFR-GFP γδ T cell (n=4). **(E)** Frequency of distinct γδ TCR subsets in αCD19-RQR8 or αCD19-DHFR-GFP γδ T cell populations at day 22 (n=4).

#### Supplementary Figure 6

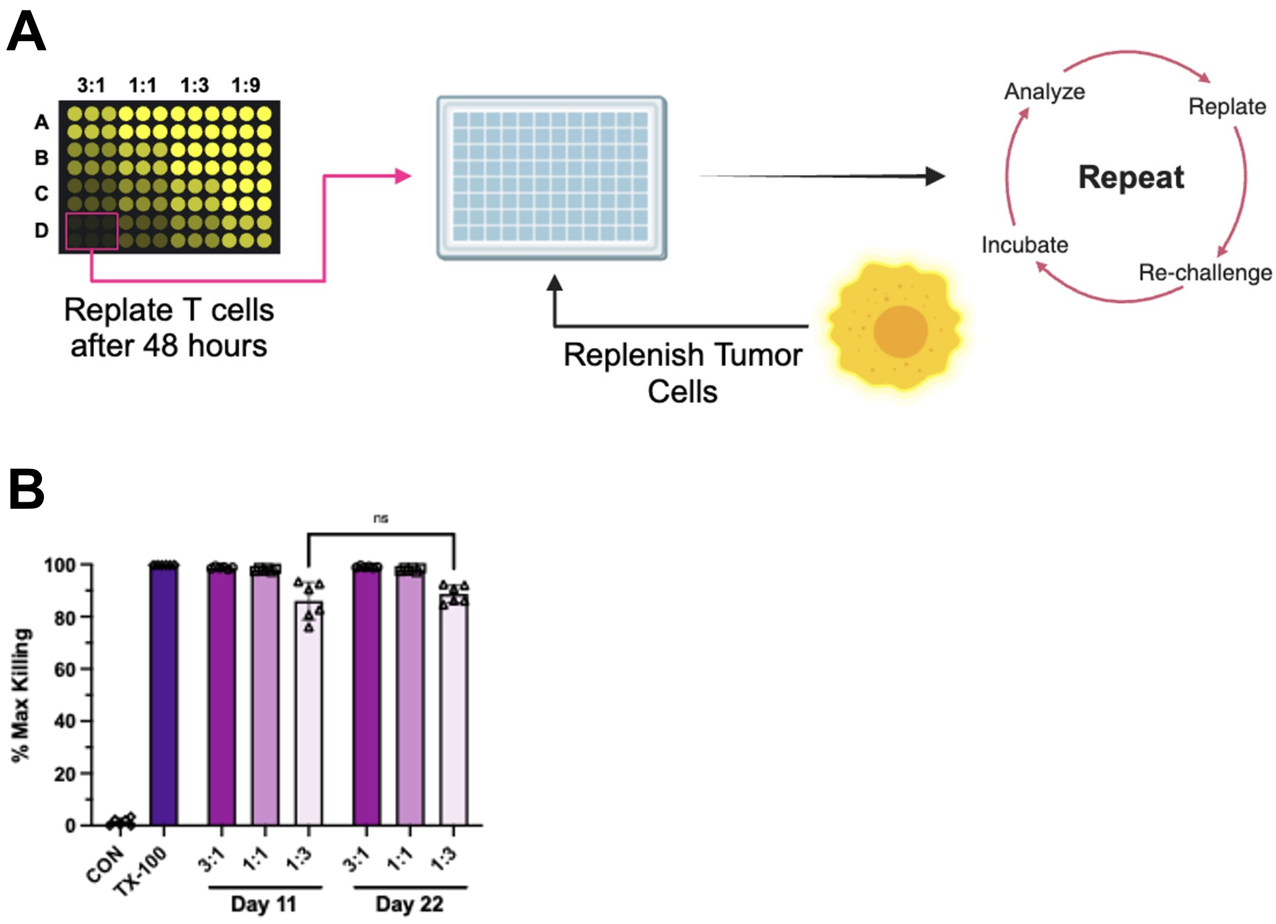


**Supplementary Figure 6. Serial killing assays.**
**(A)** Diagram of a serial killing assay. CD19-CAR γδ T cells were plated at the indicated E:T ratio with Raji-Luc cells. The cells were then replated at the same E:T ratio with fresh Raji-Luc cells after 48 hours for a total of 6 rounds of coculture (n=6). **(B)** Relative cytotoxicity of CAR-γδ T cells harvested at days 11 and 22. Cells were engineered with αCD19-DHFR-GFP at day 2, then selected with methotrexate between days 5 and 11. Percent killing was determined as a ratio of luminosity compared with Raji-Luc cells-only negative control wells and 1% Triton X-100 (TX-100) treated positive control wells. (n=2 donors with 3 technical replicates).

#### Supplementary Figure 7

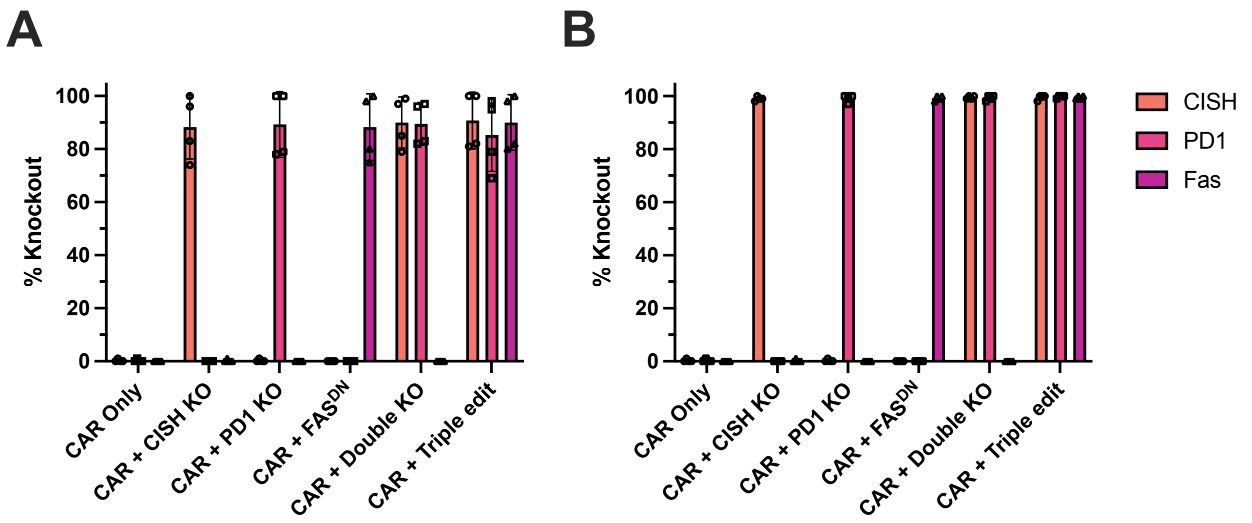


**Supplementary Figure 7. Gene editing.**

Gene editing efficiency in **(A)** unselected and **(B)** MTX-treated CAR-γδ T cells that also received a αCD19-DHFR-GFP construct on day 2. Percent knockout was assessed by DNA sequencing on day 22 (n=3).

**Supplementary Figure 8**

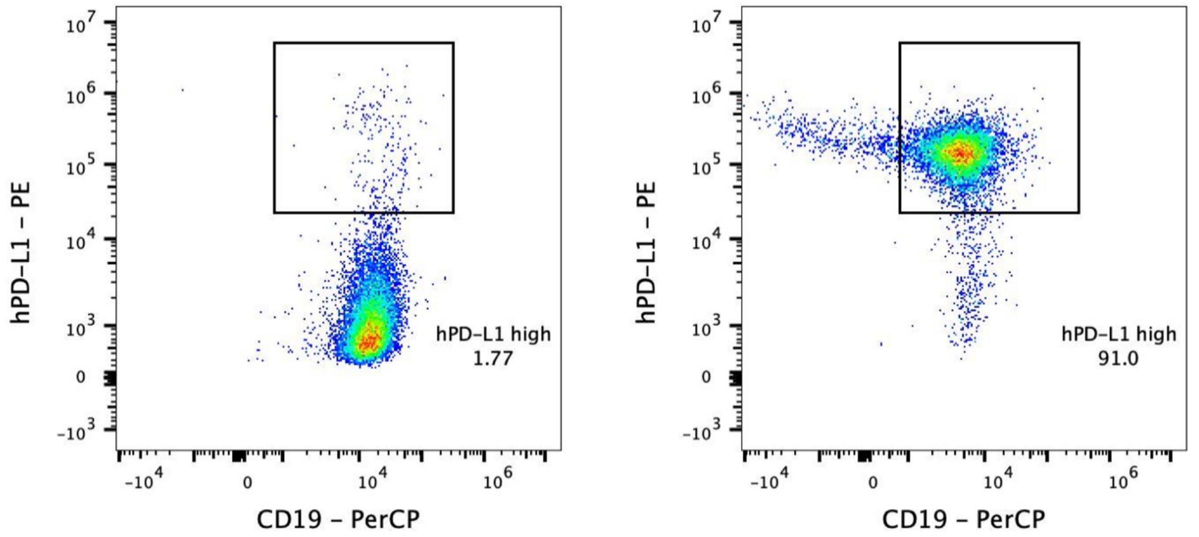


**Supplementary Figure 8. Expression of PDL1 on target cell lines.**
Expression of PDL1 in PDL1- and PDL1+ Raji cells as measured by flow cytometry.

### Supplementary Figure 9

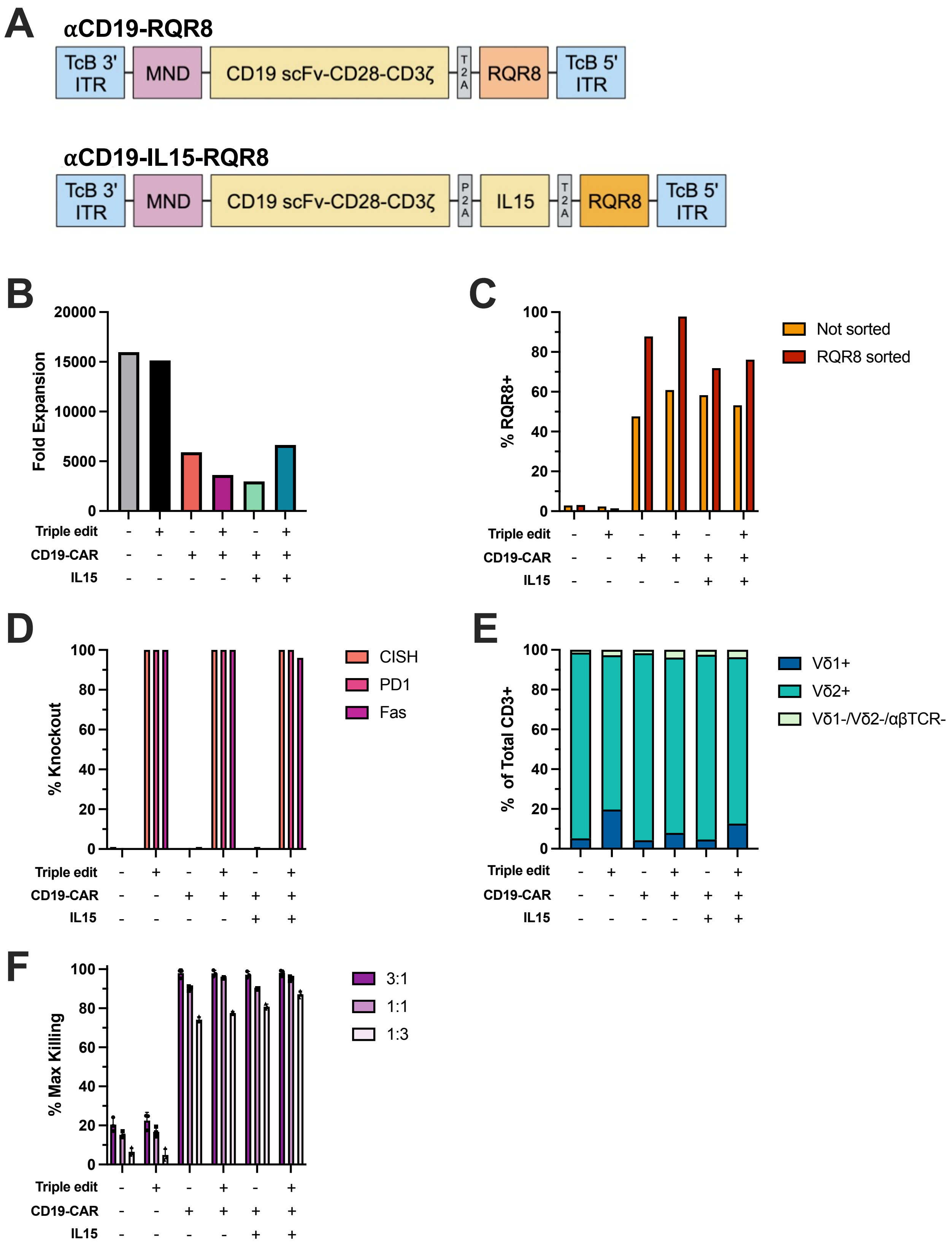

**Supplementary Figure 9. Characterization of γδ T cells used for *in vivo* challenge.**
**(A)** Constructs used to manufacture cells for in vivo challenge. **(B)** *Ex vivo* fold expansion of the infusion products. Six conditions were tested. 1) Pulse control, 2) Pulse with *PDCD1 KO*, *CISH* KO and $FAS^{DN}$, 3) αCD19-CAR, 4) αCD19-CAR with IL15, 5) αCD19-CAR, *PDCD1 KO*, *CISH* KO and $FAS^{DN}$, and 6) αCD19-CAR with IL15, *PDCD1 KO*, *CISH* KO and $FAS^{DN}$. **(C)** RQR8 expression in expanded D22 cells before and after RQR8 enrichment using immunomagnetic sorting with an anti-CD34 QBEND/10 antibody, as assessed by flow cytometry. **(D)** Gene editing efficiency, determined through DNA sequencing, in selected D22 cells, **(E)** relative frequency of distinct γδ TCR subsets, and **(F)** validation of γδ T cell cytotoxicity.

#### Supplementary Figure 10

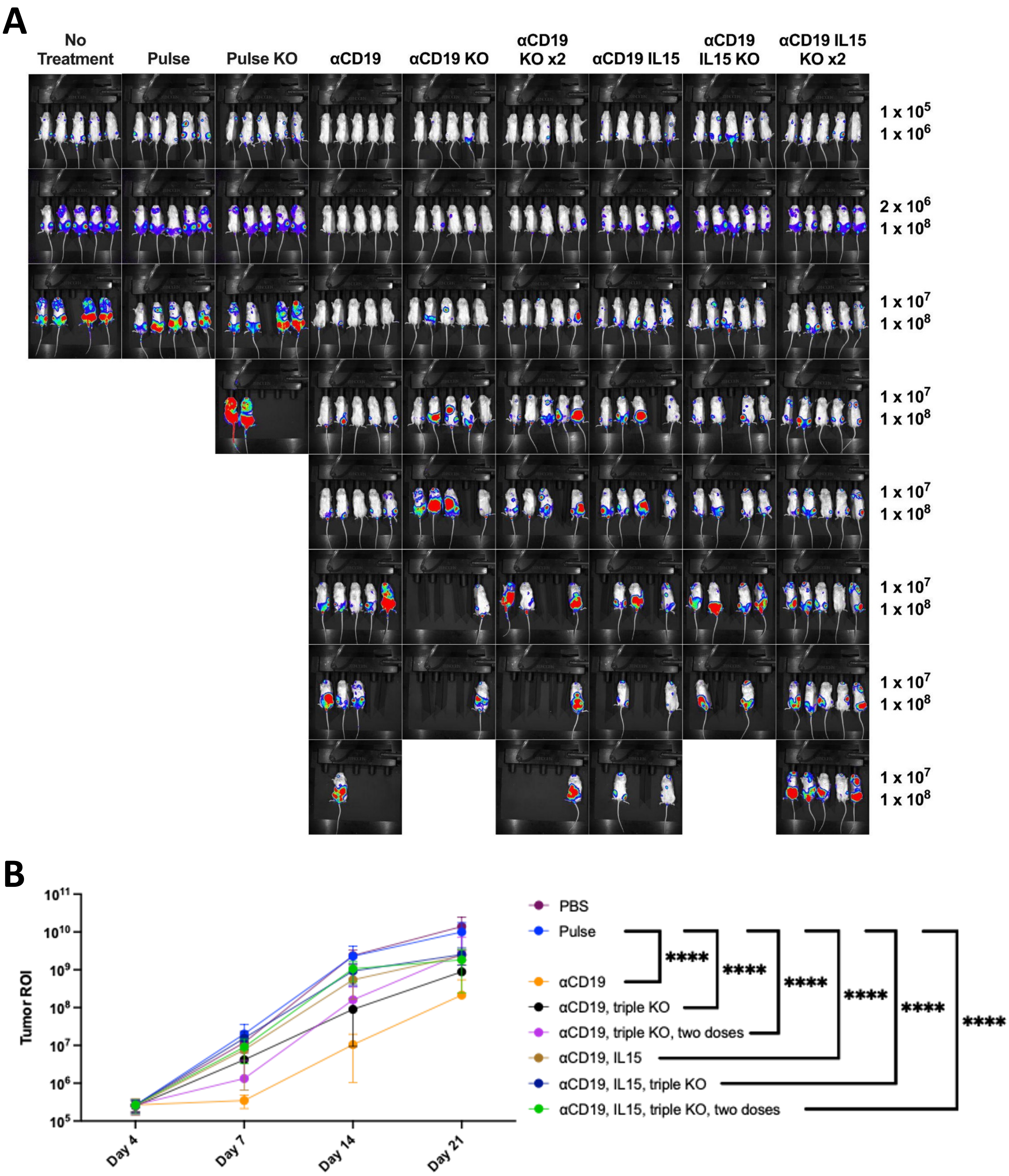


**Supplementary Figure 10. Weekly tumor fluorescence beginning at day 7 post-injection. (A)** Images were collected on an IVIS® system, then assessed in ImageJ. Minima and maxima for each week's ROI scale are listed on the right. **(B)** Statistical comparison of treatment groups to the mice that received Pulse γδ T cells for the first 21 days of the experiment, after which mouse endpoints preclude comparison. The Pulse, triple edit group was excluded from this analysis due to a missing data point. (Two way ANOVA with Dunnett's multiple comparisons test, n=5).

#### Supplementary Figure 11

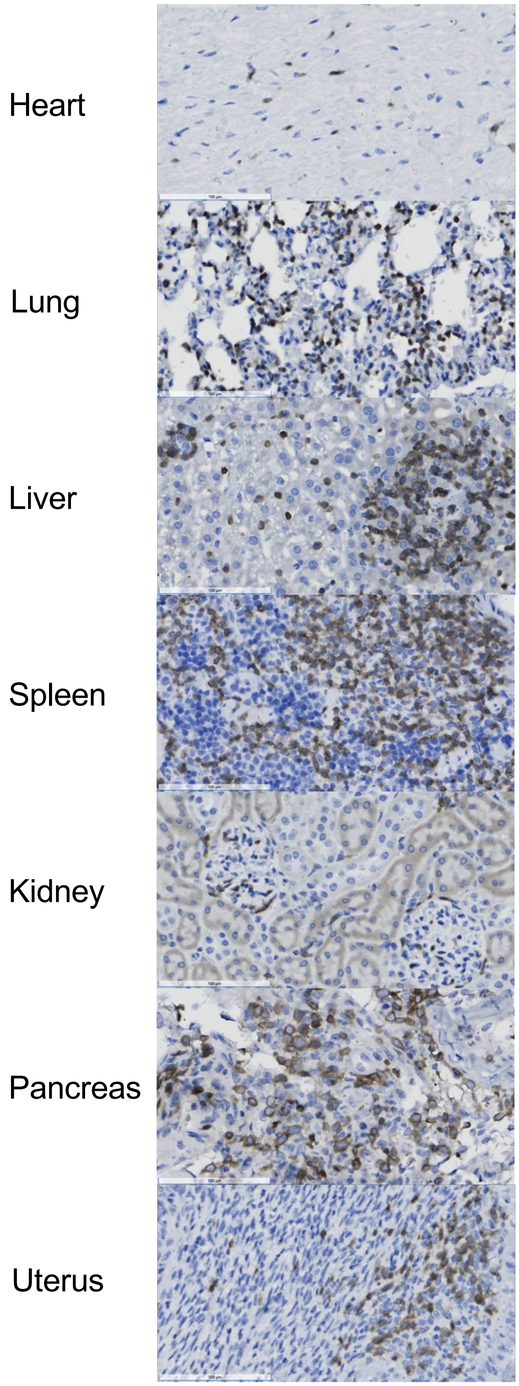


**Supplementary Figure 11. Infiltration of tissue by γδ T cells.**
Immunohistochemistry staining of CD3 in various tissues.

#### Supplementary Figure 12

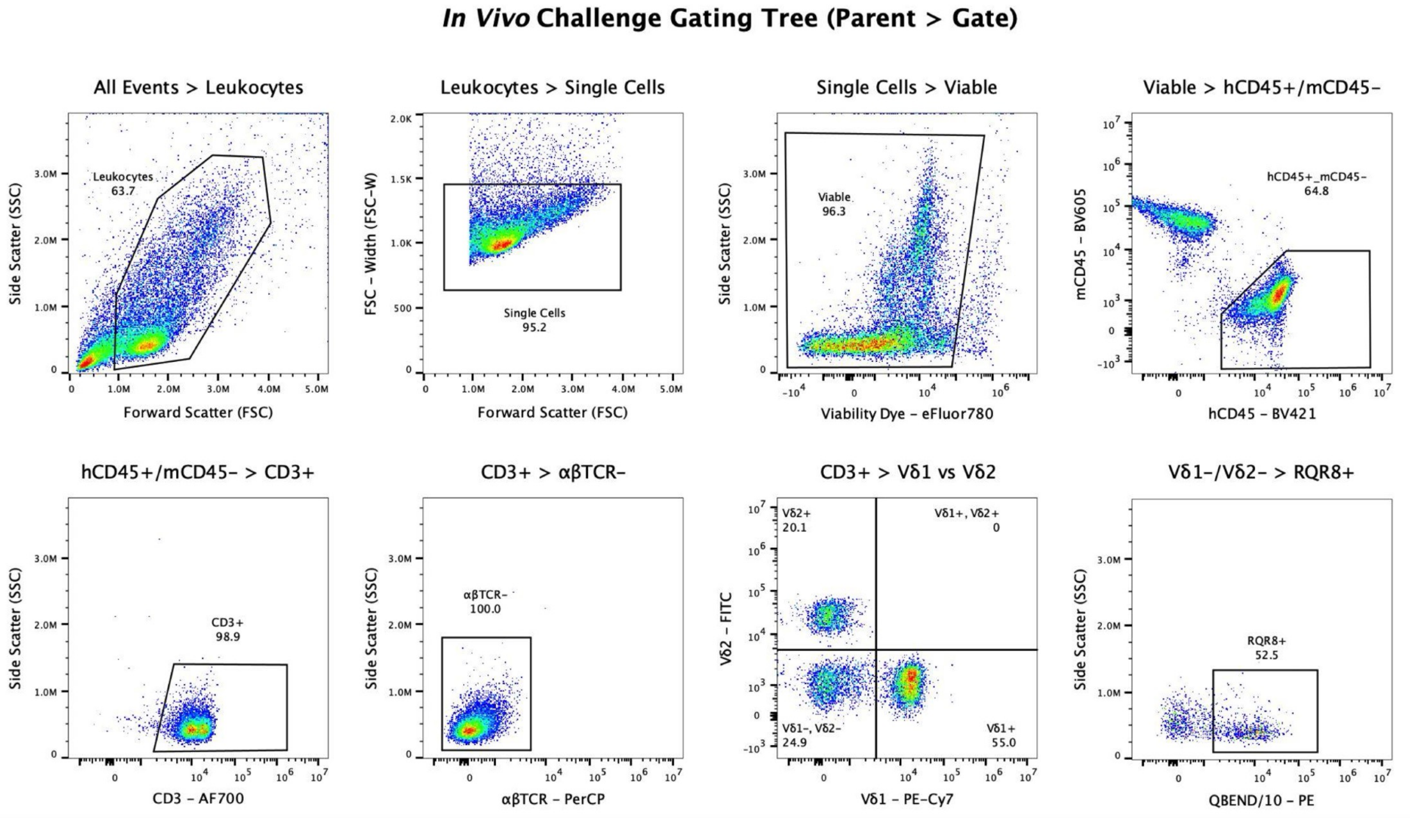


**Supplementary Figure 12. Representative flow cytometry gating tree for phenotyping γδ T cells after weekly bleeds.**

#### Supplementary Figure 13

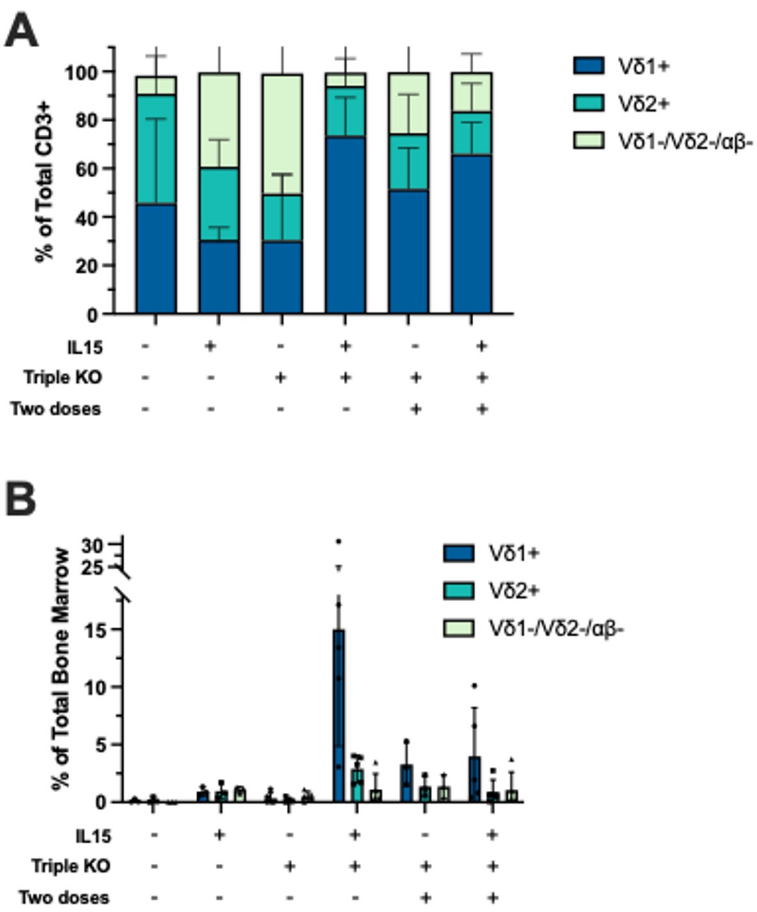


**Supplementary Figure 13. All γδ T cell subset migrate to the bone marrow.**
**(A)** Frequency γδ TCR subsets as a fraction of total CD3+ cells or **(B)** a fraction of total cells in mouse bone marrow at endpoint (n=5).

### Supplementary Figure 14

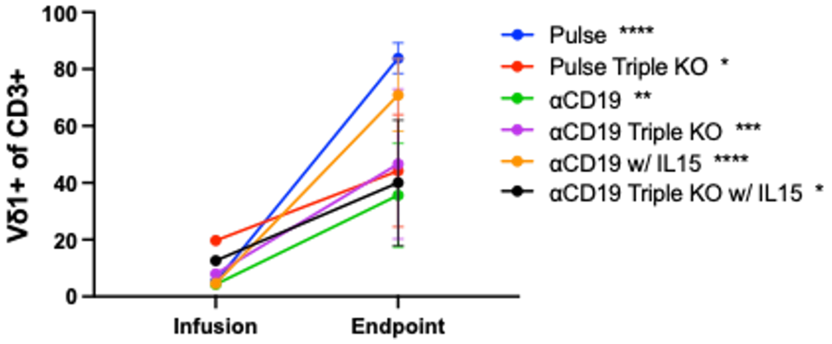


**Supplementary Figure 14. Fraction of Vδ1+ cells increases from infusion to endpoint in all conditions.**

Line graphs shows percent of Vδ1+ cells of CD3+ cells at both the infusion at endpoint for all conditions. Only mice that received a single dose of cells are included. Asterisks indicated statistically significance of changed between infusion at endpoint for each condition. Two way ANOVA with Šídák's multiple comparisons test (n=5).

#### Supplementary Figure 15

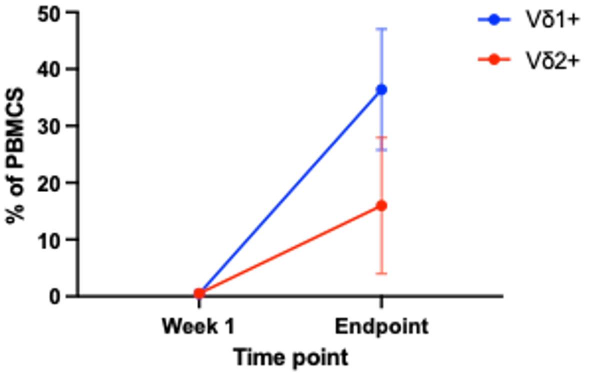


**Supplementary Figure 15. Both Vδ1+ and Vδ2+ subsets expand in the peripheral blood in mice treated with CAR γδ T cells including IL15.**

Frequency of Vδ1+ (*blue circles*) and Vδ2+ (*red circles*) γδ T cells as a percent of total PBMCs at the one week time point and at end point.

#### Supplementary Figure 16

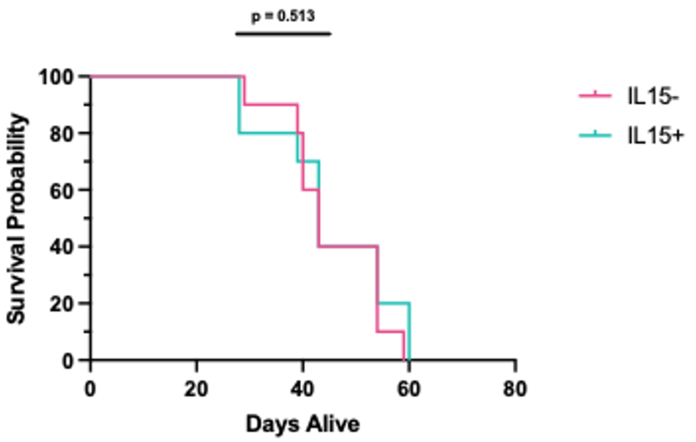


**Supplementary Figure 16. IL15 expression has no impact on survival.**
Kaplan Meier survival graphs of pooled groups of mice that received a single dose of cells engineered with constructs encoding IL15 with matched pooled groups of mice that received a single dose of cells engineered with constructs that did not code for IL15.

#### Supplementary Figure 17

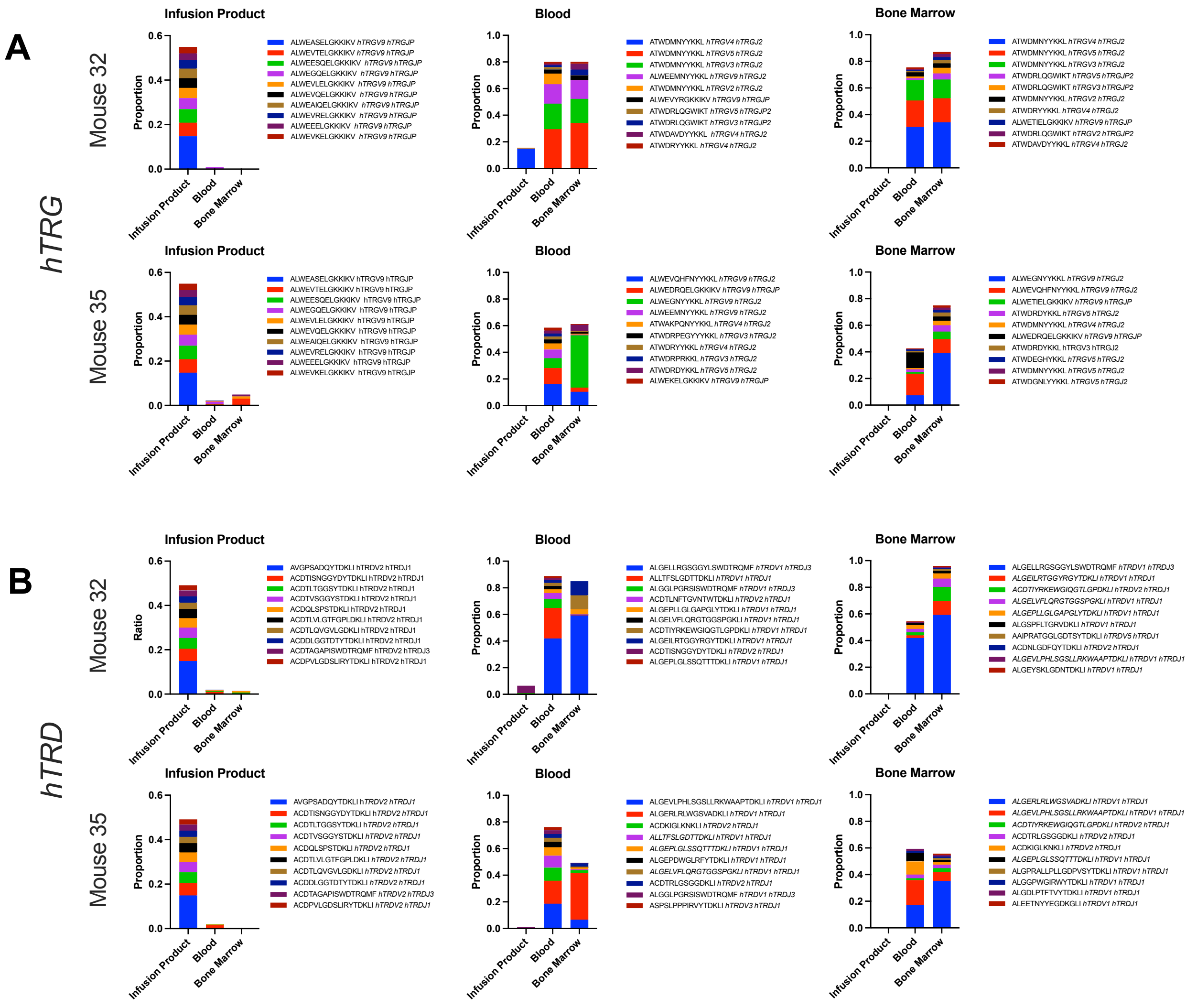


**Supplementary Figure 17. TCR clones**

Top ten most frequent **(A)** *hTRG* and **(B)** *hTRD* clones identified in the infusion product (*left column*), blood (*middle column*), and bone marrow (*right column*) compared to their frequency in other compartments.

**Supplementary Figure 18**

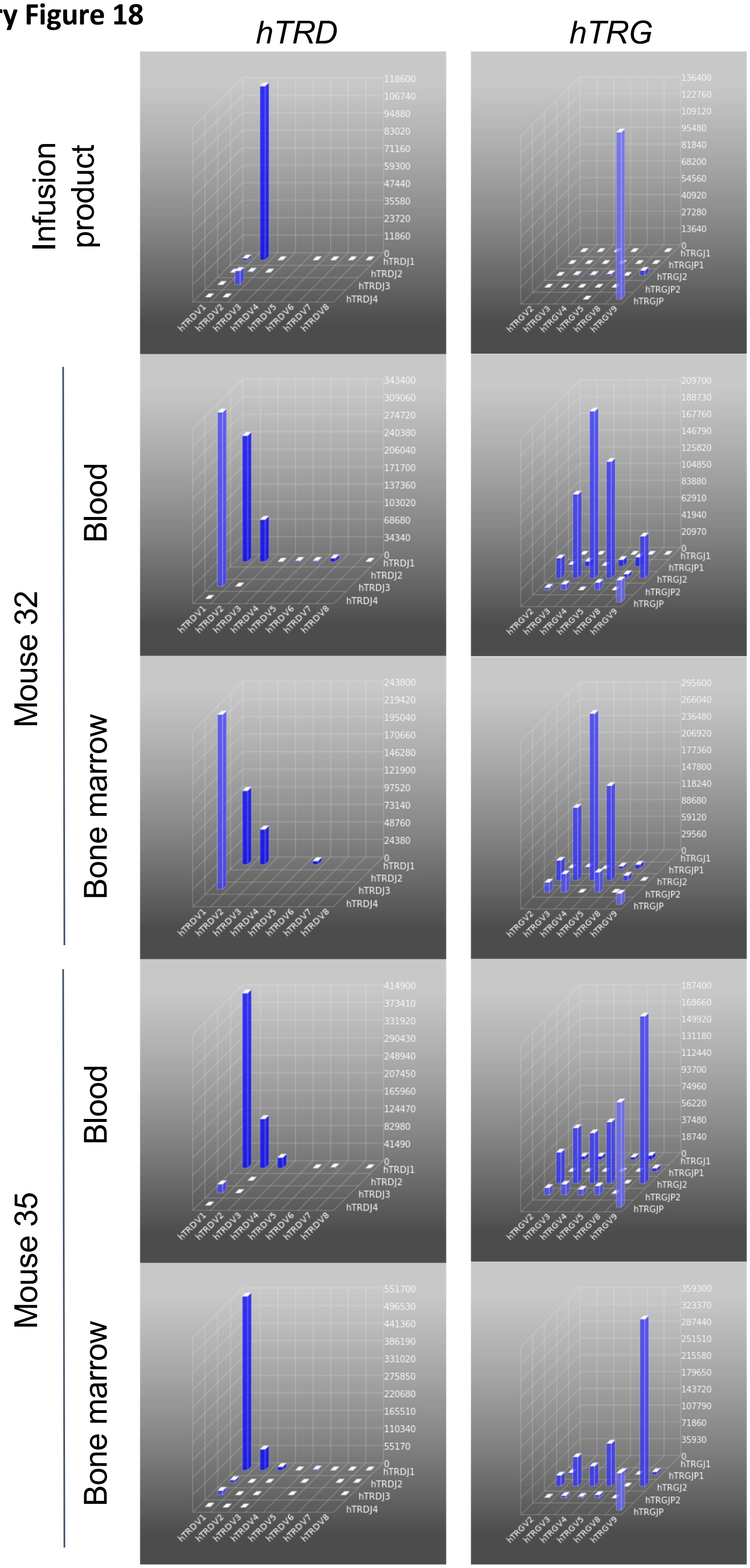


**Supplementary Figure 18. *hTRG* and *hTRD* chain diversity of the infusion product and peripheral blood of two mice at endpoint.**

#### Supplementary Figure 19

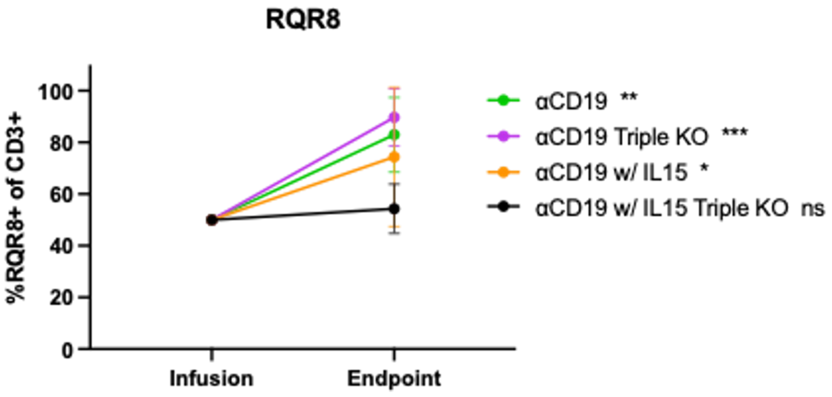


**Supplementary Figure 19. Fraction of CD19-CAR+ cell changes from infusion to endpoint.**
Line graph shows percent of CD19-CAR cells, as measured by RQR8 expression, of CD3+ cells at both the infusion and endpoint for all conditions. Only mice that received a single dose of CD19-CAR+ cells are included. Asterisks indicate the statistically significance of change between infusion product and endpoint for each condition. Two-way ANOVA with Šídák's multiple comparisons test (n=5).

#### Supplementary Table 2

| Gene | Sequence |
| --- | --- |
| *CISH* | 5'-CTCACCAGATTCCCGAAGGT-3' |
| *PD1* | 5'-CACCTACCTAAGAACCATCC-3' |
| *Fas* | 5'-AAATATATCACCACTATTGC-3' |

**Supplementary Table 2. gRNA sequences for γδ T cell functional edits.**

### Supplementary Table 2

| Target | Fluorophore | Clone | Company | Cat# |
|---|---|---|---|---|
| CD3 | AF700 | SK7 | BioLegend | 344822 |
| CD3 | FITC | SK7 | BioLegend | 344804 |
| CD3 | PECy5.5 | SK7 | eBiosciences | 344866 |
| Vδ2 | PE | B6 | BioLegend | 331408 |
| Vδ2 | FITC | B6 | BioLegend | 331406 |
| Vδ1 | PE-Vio770 | REA173 | Miltenyi | 130-117-697 |
| Pan-αβTCR | BV421 | IP26 | BioLegend | 306722 |
| CD56 | PE | 5.1H11 | BioLegend | 362508 |
| CD56 | APC | 5.1H11 | BioLegend | 362504 |
| CD56 | BV650 | 5.1H11 | BioLegend | 362532 |
| CD19 | PerCP | HIB19 | BioLegend | 302228 |
| CD33 | BV605 | P67.6 | BioLegend | 366612 |
| PD1 | BV605 | EH12.2H7 | BioLegend | 329924 |
| PD1 | AF647 | EH12.2H7 | BioLegend | 329910 |
| TIGIT | KIRAVIA Blue520 | A15153G | BioLegend | 372731 |
| TIM | AF700 | F38-2E2 | eBiosciences | 56-3109-42 |
| CD45RA | BV650 | HI100 | BioLegend | 304136 |
| CD45RO | BV510 | UCHL1 | BioLegend | 304245 |
| CD27 | AF700 | M-T271 | BD Biosciences | 560611 |
| CD45 | BV421 | 2D1 | BioLegend | 368522 |
| mCD45 | BV605 | 30-F11 | BioLegend | 103140 |
| CD27 | BV510 | L128 | BD Biosciences | 563092 |
| NKG2A | PE | S19004C | BioLegend | 375104 |
| NKG2C | BV605 | 134591 | BD Biosciences | 748166 |
| NKG2D | APC | 1D11 | BioLegend | 320808 |
| CD62L | AF647 | DREG-56 | BioLegend | 304818 |
| CD16 | PerCP | 3G8 | BioLegend | 302030 |
| iNKT | APC | 6B11 | Miltenyi | 130-094-839 |
| CD25 | BV650 | BC96 | BioLegend | 302633 |
| 4-1BB | PE | 4B4-1 | BioLegend | 309804 |
| CD34 | PE | QBEND/10 | ThermoFisher | MA1-10205 |

**Supplementary Table 2. Antibodies used for flow cytometry.**

#### Supplementary Table 3

| Primer Name | Sequence |
|---|---|
| CISH_Fwd | 5'-TGATGACAAGTGGGGAACGA-3' |
| CISH_Rev | 5'-TCCTGAGTAGTCTGGTGGGA-3' |
| PD1_Fwd | 5'-CAGCACTGCCTCTGTCACTC-3' |
| PD1_Rev | 5'-CTGAGCTGTGTGGCTTTGGG-3' |
| Fas_Fwd | 5'-TTCCCCTAGTCAGCTCTTCA-3' |
| Fas_Rev | 5'-CCAAGCTTTGGATTTCATTTC-3' |

**Supplementary Table 3. Primers used to validate gene-editing efficiency.**

#### Supplementary Table 4

| Primer Name | Sequence |
|---|---|
| WPRE.1_Fwd | 5'-GCTGCTTTAATGCCTCTGTATC-3' |
| WPRE.1_Rev | 5'-GGGCCACAACTCCTCATAAA-3' |
| WPRE.1_Probe | 5'-/56-FAM/TATTGCTTC/ZEN/CCGTACGGCTTTCGT/3IABkFQ/-3' |
| RNAseP_Fwd | 5'-AGATTTGGACCTGCGAGCG-3' |
| RNAseP_Rev | 5'-GAGCGGCTGTCTCCACAAGT-3' |
| RNAseP_Probe | 5'-/5HEX/TTCTGACCT/ZEN/GAAGGCTCTGCGCG/3IABkFQ/-3' |

**Supplementary Table 4. Primers used for ddPCR.**
